# Spatial and feature-selective attention have distinct effects on population-level tuning

**DOI:** 10.1101/530352

**Authors:** Erin Goddard, Thomas A. Carlson, Alexandra Woolgar

**Affiliations:** McGill Vision Research Group, McGill University, Montreal, QC, H3G1A4, Canada; ARC Centre of Excellence in Cognition and its Disorders (CCD), Macquarie University, Sydney, NSW, 2109, Australia; Perception in Action Research Centre (PARC) and Department of Cognitive Science, Macquarie University, Sydney, NSW, 2109, Australia; School of Psychology, University of Sydney, Sydney, NSW, 2006, Australia

**Author notes:** Corresponding Author: Erin Goddard, McGill Vision Research Group, 1650 Cedar Ave., Rm L11 513, Montreal, QC, H3G1A4, Canada. Conflict of interest: None to report.

## Abstract

Selective attention is fundamental to cognitive activity and can be deployed in different ways. Non-human primate data suggests that spatial and feature-based visual attention have qualitatively different effects on neural tuning, but this has been challenging to assess in humans. Using multivariate decoding of MEG data, we tracked the effects of spatial and feature-selective attention on population-level coding of novel objects. We found that spatial and feature-selective attention interacted multiplicatively to enhance object representation. Moreover, the two types of attention induced qualitatively different patterns of enhancement in occipital cortex, and these differences were accounted for by the principles of response-gain and tuning curve sharpening derived from single-unit work. A novel information flow analysis further showed that stimulus representations in occipital cortex were Granger-caused by coding in frontal cortices earlier in time. We find that human spatial and feature-selective attention rely on qualitatively different, interacting, neural mechanisms.

---

At any moment, there is far more information available from our senses than we can possibly process at once. Accordingly, only a subset of the available information is processed to a high level, making it crucial that brain can dynamically devote greatest processing resources to the most relevant information. Our ability to selectively attend to relevant information is remarkably flexible. For instance, we can adapt our attentional state by directing our attention in space (spatial attention, e.g. attend left), to a specific feature dimension (feature-*selective* attention, e.g. detect changes in color across a scene) or based on a particular feature value along that feature dimension (feature-*based* attention, e.g. find all the red objects), using the definitions of Chen et al. (2012). Each of these types of attention can change behavior, improving performance related to the attended location or feature-dimension, while decreasing performance on the ignored dimension/location (Pestilli and Carrasco, 2005; Rossi and Paradiso, 1995; Saenz et al., 2003; Carrasco, 2011), consistent with neural resources being redistributed.

What is the neural basis for this important ability, and to what extent do the same mechanisms give rise to spatial and feature-based attentional enhancements? Shifts in attention induce changes in the responses of individual neurons (Sprague et al., 2015; Reynolds and Heeger, 2009; Maunsell, 2015), change the overall responsiveness of brain regions (Corbetta et al., 1990; Chawla et al., 1999; Saenz et al., 2002, 2003; Serences and Boynton, 2007; Gouws et al., 2014), and change the information carried by a population response (Guggenmos et al., 2015; Woolgar et al., 2015; Vaziri-Pashkam and Xu, 2017). The most marked difference between spatial and feature-based attention is that the effects of spatial attention vary according to the part of the visual field to which a cell responds, whereas feature-based attention is spatially diffuse, changing the responses of neurons (Treue and Martinez-Trujillo, 1999; McAdams and Maunsell, 2000; Martinez-Trujillo and Treue, 2004) and voxels (Saenz et al., 2002; Serences and Boynton, 2007) across the visual field, rather than being restricted to the attended location or the stimulus location.

The reported effects of spatial attention on the tuning of individual neurons are diverse: its effects have been characterized as multiplicative response gain (McAdams and Maunsell, 1999; Treue and Martinez-Trujillo, 1999; Lee and Maunsell, 2010b), contrast gain (Li and Basso, 2008; Martinez-Trujillo and Treue, 2002; Reynolds et al., 2000), or a combination of these effects (Williford and Maunsell, 2006). There have also been mixed results regarding the effect of spatial attention on contrast response functions measured with fMRI (Buracas and Boynton, 2007; Li et al., 2008). Fewer studies have investigated the effects of feature-based attention, and only a subset of these where shifts in feature-based attention were not accompanied by changes in spatial attention (Maunsell and Treue, 2006). Intriguingly, feature-based attention may affect the tuning of individual neurons in a subtly different manner to spatial attention. In an influential electrophysiological study Martinez-Trujillo and Treue (2004) found effects at the single-unit level which would lead to a ‘sharpening’ of the population response around the attended feature value across the visual field. In a recent MEG study Bartsch et al. (2017) reports similar sharpening of the population response with attention to color. However, even this difference in the effects of spatial and feature-based attention does not eliminate the possibility of a unified attentional system, where stimulus location is treated as one of many stimulus features that can potentially be selected with attention (Treue and Martinez-Trujillo, 1999; Maunsell and Treue, 2006; Maunsell, 2015).

While there is an increasing body of work investigating the effects of spatial and feature-based or feature-selective attention, there are few studies that directly compare these two attention types. In one of the few previous studies that simultaneously manipulated both spatial and feature-selective attention Cohen and Maunsell (2011) implied highly similar processes of spatial and feature-selective attention, affecting the same subpopulations of neurons. The main difference between their effects was across hemispheres: for feature-selective but not spatial attention the effects were correlated across hemispheres. Directly comparing attention types is critical for resolving whether and how these attention types produce different effects on the population code, and how their effects interact.

The overlapping characteristics of these two attention types make them difficult to separate, as does the diversity of their reported effects. Another complicating factor is that much of our current understanding of attention comes from work exploring its effects on individual neurons, but attention can also induce changes in the information represented by a population of neurons that will not be revealed in the tuning curves of individual neurons (Sprague et al., 2015). For instance, attention has been shown to decrease response variance (e.g. Mitchell et al. 2007), and decrease (Cohen and Maunsell, 2009) or increase (Ruff and Cohen, 2014) the correlation between pairs of neurons. It can be difficult to predict how each of changes should affect the information represented by the population response, for example, predicting how changes in correlation across neurons will affect population codes is non-trivial (Moreno-Bote et al., 2014). There is a need, then, to complement measurements of the effects of attention on single-unit responses with measurements of its effects on information carried by a population of cells (Sprague et al., 2015), via simultaneous multi-electrode recordings (Cohen and Maunsell, 2011) or neuroimaging. Multivariate classification analyses, applied to multi-electrode recordings or neuroimaging measures, provide a means of measuring the overall stimulus-related information that is carried by a population response. Unlike the tuning of single neurons, any signal or noise correlations that could decrease or increase information carried by the population response (Moreno-Bote et al., 2014) should affect classifier accuracy. This sensitivity to additional factors make classifier accuracy an ideal intermediate level of description for linking single-unit responses to the information in the population response which is available for readout by other brain regions, and to the organism’s percept/behaviour (Carlson et al., 2018).

Another key question for understanding attentional modulation of visual information is to identify the regions that drive these changes in processing, and when and how they influence visual cortical areas. There is evidence that some prefrontal cortical (PFC) regions are critically involved in visual attention and task-based modulations in response, including the frontal eye fields (Moore et al., 2003; Gregoriou et al., 2012; Zhou and Desimone, 2011), the ventral prearcuate region (Bichot et al., 2015), the superior precentral sulcus (Jerde et al., 2012) and lateral PFC (Tremblay et al., 2015; Luo and Maunsell, 2018). Selective prioritisation of task-relevant information in prefrontal cortex (e.g. Duncan 2001) may provide a source of bias, driving processing in visual cortices in favour of task relevant information (Desimone and Duncan, 1995; Dehaene et al., 1998; Miller and Cohen, 2001). But precisely what this influence is, and when it occurs, remains unknown.

Here we measured the effects of spatial and feature-selective attention within the same datasets of magnetoencephalography (MEG) recordings (n=20), enabling us to directly compare and contrast their effects. We obtained fine timescale measures of stimulus-related information in two large regions of interest (ROIs): visual cortex and frontal/prefrontal cortex. For both ROIs, we found strong, multiplicative effects of spatial and feature-selective attention, but these only emerged relatively late (*>*200ms after stimulus onset). We used an information flow analysis to test for how the two ROIs were interacting over time: we measured Granger-causal relationships between their stimulus-related information. This revealed that for visual cortex, the strongest attentional modulation occurred after the onset of feedback from frontal regions. We also tested whether spatial and feature-selective attention induced different effects on the population response. We predicted that both types of attention would enhance stimulus-related information, but that feature-selective attention would induce sharpening of the population response around the attended feature value, whereas spatial attention would induce a more generalized enhancement across feature values. In line with these predictions, we found that spatial attention produced relatively more enhancement of discriminability for stimulus pairs that were far apart in feature space, while the effects of feature-selective attention were relatively stronger for stimulus pairs that were closer in feature space.

## Results

### Performance on behavioral task

Participants (n=20) viewed a series of stimuli while we recorded their neural activity using MEG. On every trial there were two objects on the screen, one on the left and one on the right of fixation (Figure 1**A**). Participants were instructed to covertly attended either to the stimulus on the left or right of fixation (spatial attention manipulation), and they were required to make a judgment based on the target object’s color or shape (feature-selective attention manipulation). As shown in Figure 1**B**, there were four stimulus colors ranging from red to green, and four shapes ranging from strongly X-shaped to strongly non-X-shaped. The four feature values along each dimension meant that for both tasks the stimuli were either far from the decision boundary (e.g. strongly red; ‘easy’ trials) or closer to the decision boundary (e.g. weakly red; ‘hard’ trials). As expected, participants were faster and more accurate at identifying color and shape for objects that were far from the decision boundary relative to those that were near the decision boundary. For the color task, the average accuracy was 95.6% (std 3.6%) on the easy trials, and 85.2% (std 7.3%) on the hard trials, while median reaction time was 0.69s on the easy trials and 0.81s on the hard trials. Similarly, for the shape task the average accuracy was 94.1% (std 3.5%) on the easy trials, and 74.1% (std 4.7%) on the hard trials, while median reaction time was 0.74s and 0.82s on the easy and hard trials respectively.

**Figure 1:**
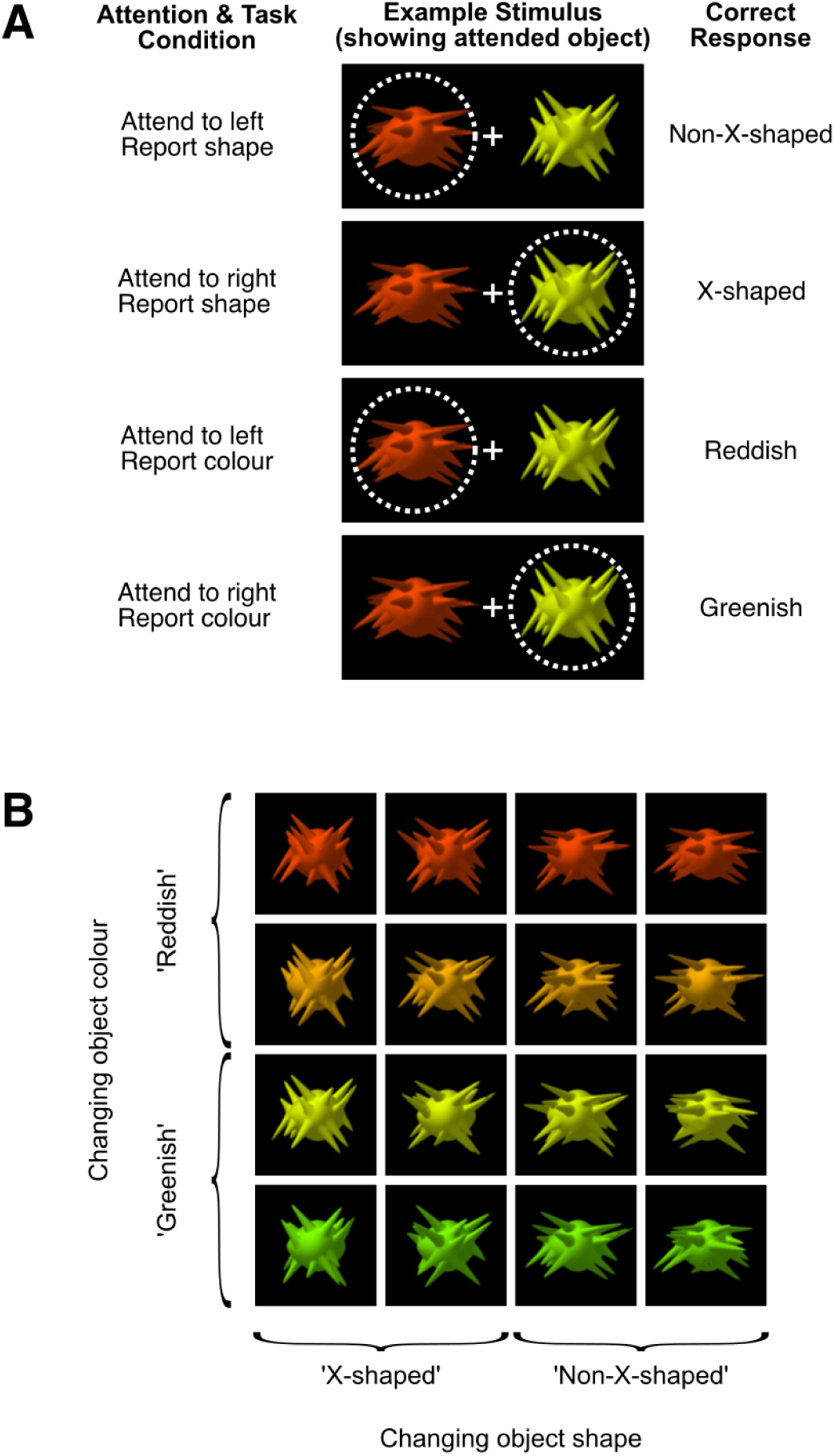
Visual stimuli, showing attention conditions (**A**) and stimulus dimensions (**B**). **Attention conditions (A):** At the start of each block of trials, participants were told the location to which they should direct their attention (left or right of fixation), and which task they should perform for that block of stimuli: either reporting on the target object’s shape (‘X-shaped’ or ‘non-X-shaped’) or color (reddish or greenish). Two objects appeared on each trial, and participants covertly attended to one while we used eye tracking to monitor their fixation. The example above illustrates how the same stimulus configuration was used in each of the four attention/task conditions. The dotted circle indicates the location of spatial attention, and was not visible during the experiment. **Stimulus dimensions (B):** Each object varies systematically along 2 dimensions, color and shape. In the color task, participants categorized the attended object as either ‘greenish’ or ‘reddish’. In the shape task, participants categorized the attended object as either ‘X-shaped’ or ‘non-X-shaped’, based on the orientations of the object’s spikes. To encourage participants to attend to the overall object shape rather than (for example) the orientation of a single spike, on each trial the object was randomly selected from 100 exemplars with the target shape statistics, and there were variations between exemplars in the location, length and orientation of the spikes. This is illustrated above in the shape variation between objects in the the same column.

### Decoding attentional state

We trained classifiers to make a series of orthogonal discriminations in order to quantify neural information about the participant’s task and the stimulus. First, we trained classifiers to discriminate the participant’s attentional set: the attended location (left versus right) and feature (color versus shape). Second, we trained classifiers to discriminate the stimuli and compared the strength of discrimination between attentional conditions.

Our first question concerned the timecourse with which we could decode information about the participant’s attentional state. For both ROIs we asked whether we could decode where participants were attending (left or right) and what task they were performing (color or shape) at each timepoint. Figure 2 shows that attentional state could be decoded from both occipital and frontal sources at most time points (at most time points the between-subjects mean was above zero when tested with a one-tailed *t*-test, *p* < 0.05, FDR corrected at *q* < 0.05 for multiple comparisons across time points, (Genovese et al., 2002)). The period of above-chance classifier performance for attended location included time points before the onset of the stimulus, when participants knew their task and were waiting for the stimulus to appear: classifier performance at this time was low but significantly above chance for both ROIs. Although we do not have a behavioral measure of the participant’s attentional state at this time, these pre-stimulus effects suggest that neural activity differed with the location to which participants were covertly attending, or to which they were preparing to covertly attend. This interpretation is consistent with previous work demonstrating the pre-stimulus effects of spatial attention on neural coding (Kastner et al., 1999; Ress et al., 2000).

**Figure 2:**
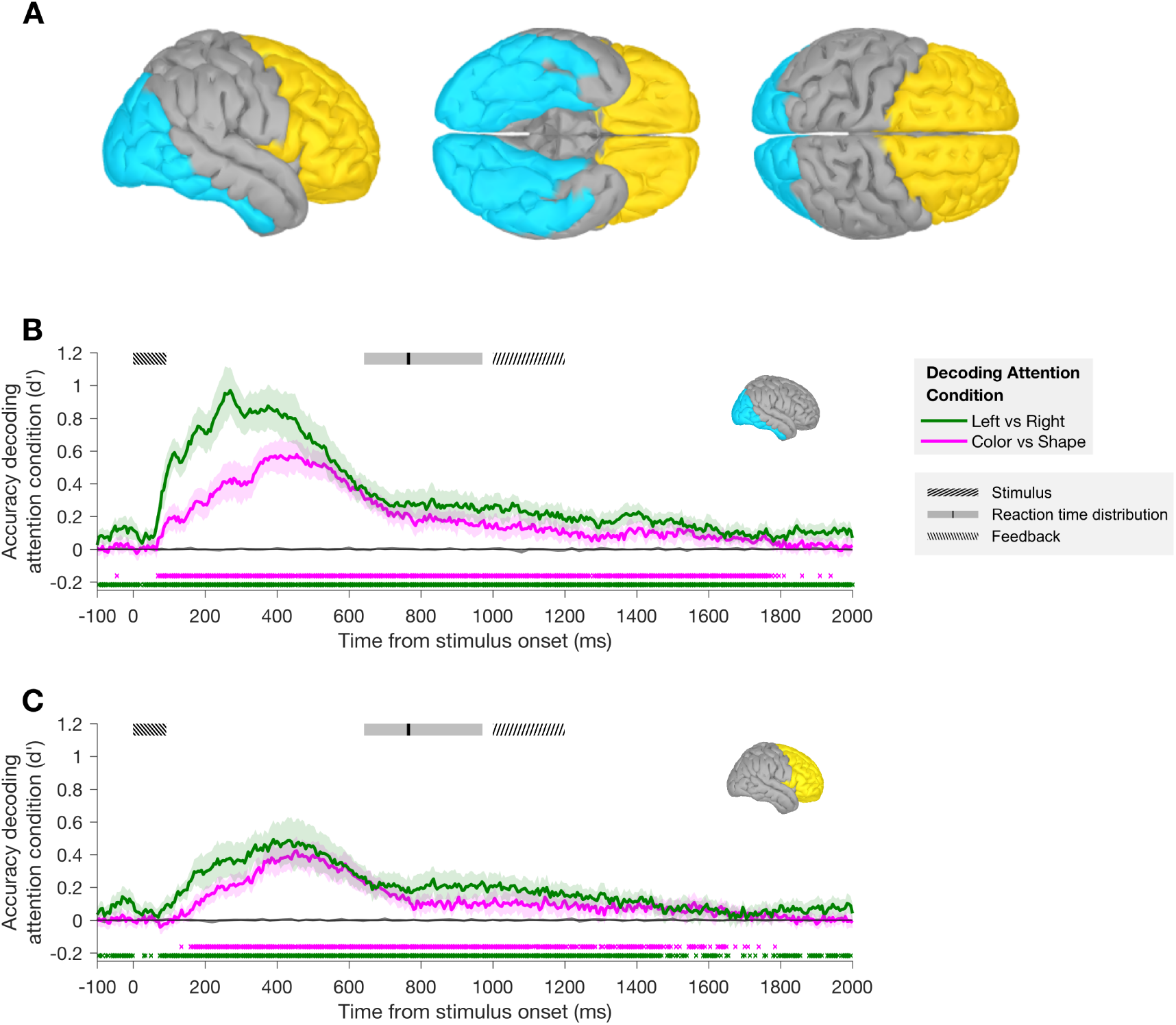
Regions of interest (A) and classifier performance across participants (n=20) for decoding attention condition using occipital sources (B) or frontal sources (C). **A**: the ‘Occipital’ (cyan) and ‘Frontal’ (yellow) regions of interest shown on the partially inflated cortical surface of the ICBM152 template brain. **B** and **C**: At each timepoint, classifiers were trained to discriminate the location and feature to which participants were attending. The shaded error bars indicate the 95% confidence interval of the between-subject mean. At the top of each plot, boxes indicate the time of the stimulus presentation (shaded area indicates onset until the median duration of 92ms), the reaction time (RT) distribution (shaded area includes RTs within the first and third quartiles, black line indicates median RT), and the time during which participants received feedback on their accuracy on those trials where their RT was *<*1s (77% of trials). On trials where RT was *>*1s (23% of trials), the 200 ms feedback started at the time of response. Classification performance can be above chance in the pre-stimulus period since attentional condition was blocked: participants knew which attentional condition to perform before the stimulus appeared. Nonetheless, decoding of attentional condition improved dramatically after the stimulus was presented, and peaked earlier when classifiers were decoding attended location (270 ms and 390 ms after stimulus onset for occipital and frontal ROIs respectively) than when decoding attended object feature (455 ms after stimulus onset for both ROIs). Shaded gray region around x-axis indicates the 95% confidence intervals of the same classifications when performed on permuted data (chance performance level). Colored crosses below the plot indicate that at every time point classifier accuracy was significantly above the average chance performance level (chance d’= 0.0001 (A), 0.0000 (B); *p* < 0.05 in a one-tailed *t*-test of the between-subject mean, FDR corrected at *q* < 0.05 for multiple comparisons across time points).

Decoding of both attended location and attended object feature increased substantially once the stimulus appeared. This presumably reflects changes in neural activity associated with enhancing the neural representation of the attended object and the task-relevant feature and/or suppressing the neural representation of the unattended object and the task-irrelevant feature of the attended object. Classification of attended feature was above chance from 70*ms* and 135*ms* after stimulus onset in the occipital and frontal ROIs respectively. Classifier performance peaked earlier when classifiers were decoding attended location (270*ms* and 390*ms* after stimulus onset for the occipital and frontal ROIs respectively) than when decoding attended object feature (455*ms* after stimulus onset for both the occipital and frontal ROIs). These timing differences could reflect differences between the timing of spatial and feature-selective attention processes, but these data are not conclusive (particularly for the onset times) since the lower overall accuracy for decoding of attended feature may have contributed to the delay in onset of above-chance classifier performance (Grootswagers et al., 2017).

### Decoding object features: color and shape

We next asked whether we could use the neural signal to decode the features of the attended and unattended stimuli, and how this information varied over time and attentional state. Our design included simultaneous manipulations of both attended feature and attended location, enabling us to ask how these different types of attention interact. By balancing the training trials across irrelevant features and creating averaged ‘pseudo-trials’ (see Methods), we were able to train classifiers to discriminate the color and shape of both the attended and non-attended object. Figure 3 shows the decoding of object color and shape for each attention condition, in each case averaged across 6 pairwise comparisons, and transformed classifier weights, showing the most informative locations in each ROI, are summarized in Figure S7.

**Figure 3:**
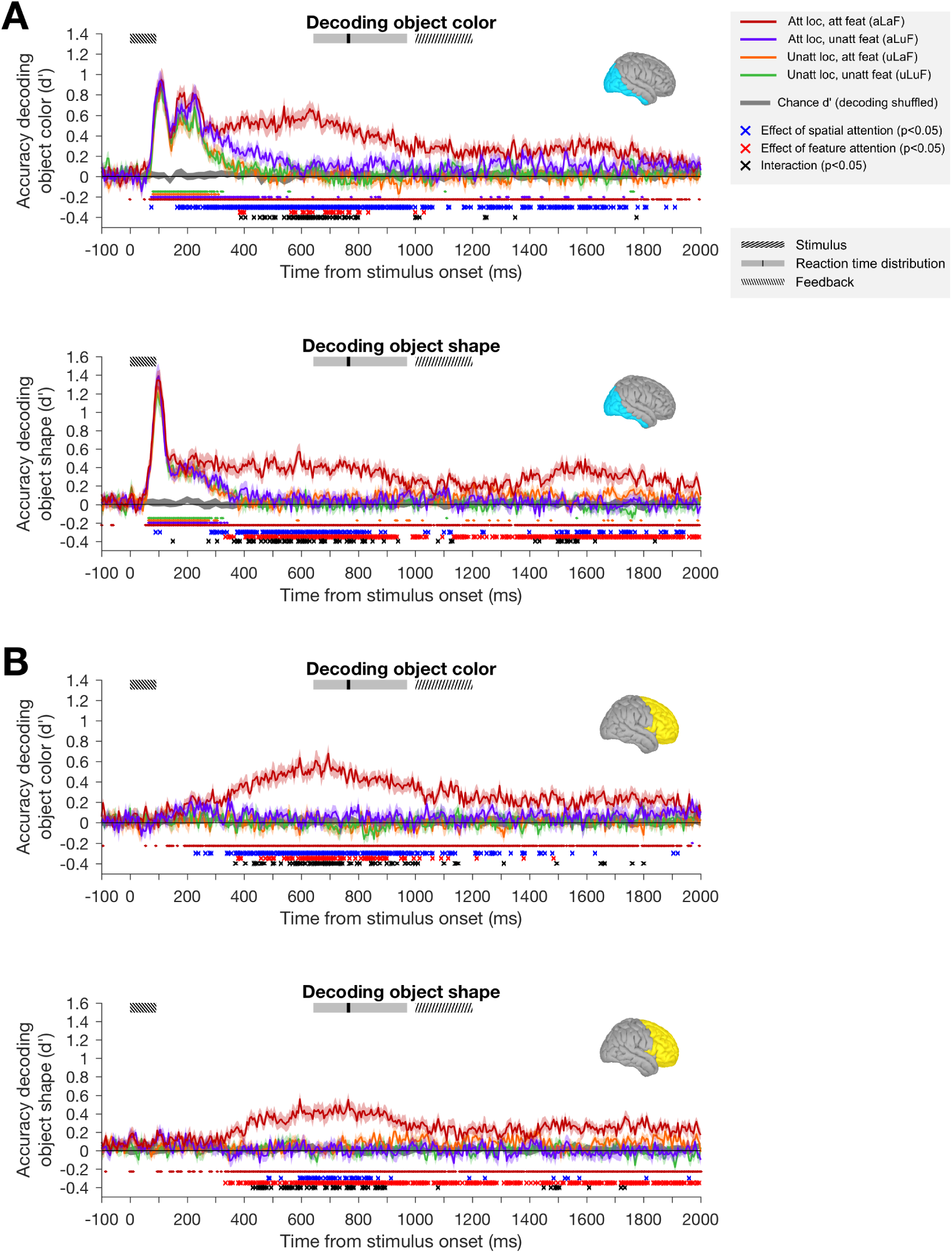
Classifier performance across participants (n=20) for decoding object features. For both occipital (**A**) and frontal (**B**) regions of interest, classifiers were trained to discriminate the color (upper plots) and shape (lower plots) of attended and unattended objects. Classifier performance is shown for each attention condition separately: attended location, attended feature (aLaF); attended location, unattended feature (aLuF); unattended location, attended feature (uLaF); and unattended location, unattended feature (uLuF). Shaded error bars indicate the 95% confidence interval of the between-subject mean, and boxes at the top of the plot show relevant trial events. The shaded gray region around the x-axis indicates the 95% confidence intervals of the four classifications when performed on randomly permuted data (the empirical null distribution). Small dots below each plot indicate timepoints at which the classification of matching color was above chance level (FDR corrected, *q* < 0.05). Below these, crosses indicate timepoints at which there was a significant effect (FDR corrected, *q* < 0.05) of spatial attention (blue asterisks), feature-selective attention (red asterisks) or an interaction of the two (black asterisks).

For both decoding object color and object shape, 2-way ANOVAs revealed significant main effects of spatial attention and feature-selective attention, and significant interactions between these effects, at the times indicated by blue, red and black crosses respectively in Figure 3 (*p* < 0.05, in each case FDR corrected at *q* < 0.05 across time points). In the occipital ROI, for both object shape and object color, we found an initial peak of robust classifier performance which showed a small effect of spatial attention, followed by a selective increase in the neural information concerning the relevant feature of the attended object, while all other information was attenuated. Around the initial peak of stimulus decoding spatial attention produced a small but significant increase in decoding of both color and shape (blue crosses < 100*ms* in Figure 3A, at 75*ms* for decoding color and 90 and 105*ms* for decoding shape). After the initial peak, the representation of task-relevant stimulus-related information was sustained, persisting beyond the offset of the stimulus (median: 92*ms*) and beyond the median response time (770*ms*). In the frontal ROI, above-chance decoding accuracy emerged later than for the occipital ROI, and was only seen for the attended feature at the attended location. This is consistent with frontal areas prioritizing representation of task-relevant information.

Interestingly, for both occipital and frontal regions, the effects of spatial and feature-selective attention interacted with each other, consistent with their effects combining in a multiplicative rather than an additive manner. For both occipital and frontal ROIs, whenever both spatial and feature-selective attention had significant effects there was generally also an interaction. The interaction reflected the selective boost in the decoding of the attended feature at the attended location, with little enhancement in classifier performance for spatial attention in the absence of feature-selective attention or for feature-selective attention in the absence of spatial attention. We think it is unlikely that the lack of an independent effect of feature-selective attention in our data reflects a true absence of any effect of feature-selective attention at the unattended location, since there are numerous reports of feature-based attention having effects at unattended locations (e.g. Treue and Martinez-Trujillo 1999; McAdams and Maunsell 2000; Martinez-Trujillo and Treue 2004; Saenz et al. 2002; Serences and Boynton 2007; Ipata et al. 2012; Bichot et al. 2015). However, there are two differences between our results and this previous work which may reflect a genuine difference. Firstly, these modulations are typically reported during responses to stimuli at the unattended location (Treue and Martinez-Trujillo, 1999; McAdams and Maunsell, 2000; Martinez-Trujillo and Treue, 2004; Saenz et al., 2002), whereas here the effects are predominantly after stimulus offset (but see Serences and Boynton 2007). Secondly, in our experiment the participants were attending to a feature dimension (feature-selective attention) rather than a particular feature value (feature-based attention), so the absence of an effect of feature-selective attention at the unattended location may reflect a difference between these types of feature attention.

Despite these protocol differences, a more parsimonious explanation is that any effects of feature-selective attention on the representation of the unattended stimulus were too small to detect. For both feature-based and feature-selective attention, a weak effect of feature attention at unattended locations is also predicted where feature attention is spatially diffuse but there is a multiplicative interaction between feature and spatial attention. The normalization model of Reynolds and Heeger (2009), which is considered in greater detail below, includes versions with either additive or multiplicative interactions between spatial and feature-based attention. The multiplicative version of their model, which is most consistent with our data, predicts a strong interaction between the effects of spatial and feature-based attention, and a very small effect of feature-selective attention alone (see Figure 7**B** and discussion below), which may have been too small to detect here. This interaction between spatial and feature-selective attention demonstrates that the neural information was highly adapted to the participant’s task, and that the brain is efficiently selecting only relevant information for sustained processing.

The earliest peaks in classifier performance for the occipital ROI showed only a slight modulation with attention. For decoding object color, the initial peak was at 105–110*ms* after stimulus onset in all attention conditions, and there was no significant effect of attended location or attended feature at either time point (2-way ANOVAs, with subject as a random factor, at 105*ms* and 110*ms*: *F*_(1,19)_ = 2.54, 2.20, *p* = .13, .15 for effect of attended location; *F*_(1,19)_ = .26, .40, *p* = .62, .54 for effect of attended feature). For decoding object shape, the initial peak was at 95–100*ms* after stimulus onset in all conditions, and there was a small increase in classifier performance at the attended location which reached significance at 95*ms* (*F*_(1,19)_ = 4.48, *p* = .048, *q* < .05 with FDR correction), and approached significance at 100*ms* (*F*_(1,19)_ = 4.36, *p* = .051). There was no significant effect of attended feature on decoding of shape at either 95*ms* or 100*ms* (*F*_(1,19)_ =.18, .41, *p* = .68, .53). The weak effects of attention on classifier performance in the occipital ROI suggest that at the time of the initial peak the object representation in visual cortex is primarily stimulus-driven. This is consistent with the lack of above-chance decoding in the frontal ROI at this time. Previous work shows that attention tends to have a greater effect on the sustained part of neural responses than on onset transients (Fries et al., 2001; Cohen and Maunsell, 2009; Lee and Maunsell, 2010a) (although the temporal dynamics of attentional modulation vary according to task requirements (Ghose and Maunsell, 2002)). The short duration of our stimulus (median: 92*ms*) means that we cannot confidently separate the sustained part of the stimulus-driven response from responses reflecting short-term memory and response preparation following stimulus offset, but our finding that the initial transient is largely unaffected by attentional task is consistent with these previous results.

There was also a secondary early peak in the occipital ROI for decoding color (∼165–240*ms* after stimulus onset), but not for decoding shape. During this second early peak for decoding color there was a significant effect of spatial attention, with stronger decoding at the attended location than at the unattended location, but classifier performance in all attention conditions remained relatively high compared to later times, where there was a marked attenuation of classification performance for all conditions except the attended feature, attended location condition.

At later time points there were stronger effects of both spatial and feature-selective attention for both stimulus features at both ROIs, and an interaction between the effects of the two types of attention. In the occipital ROI, the effect of spatial attention preceded that of feature-selective attention. For decoding object color there was a sustained effect of spatial attention from 165*ms* after stimulus onset, while the earliest significant effect of feature-selective attention was 385*ms* after stimulus onset. For shape there was a sustained effect of spatial attention from 285*ms* after stimulus onset, and an effect of feature-selective attention from 335*ms* after stimulus onset. In both cases (color and shape), the sustained effects of spatial and feature-selective attention interacted multiplicatively (seen in the selective enhancement of the aLaF condition, and the black crosses in Figure 3).

Information about the attended feature at the attended location (dark red lines in Figure 3) had later, local peaks in the vicinity of 600*ms* post-stimulus onset for both stimulus features in both ROIs: decoding of both color had local peaks at 540*ms* and 630*ms* for the occipital ROI, 595*ms* and 695*ms* for the frontal ROI; decoding of shape peaked at 590*ms* in the occipital ROI and 595*ms* in the frontal ROI. Each of these peaks are well after the offset of the stimulus (92*ms*) and just prior to the median response time (770*ms*), suggesting that classifier performance around this later peak may be associated with the participant’s decision and/or the remembered feature value. Since we balanced the response mapping (by switching the keys associated with each response pair on half the runs) it is unlikely that the motor preparation associated with the participants’ response contributed to this effect.

For both occipital and frontal ROIs, classification of the attended feature of the attended object remained above chance well after the median response time. Sustained classification of the task-relevant information could reflect processing of the feedback presented to participants after 1000*ms* (see Methods). To limit the scope of the present study we include only data from 0–1000*ms* after stimulus onset in our next analyses, excluding any effects due to the feedback.

In summary, classification performance of the occipital ROI contain early peaks in the decoding of both color and shape that showed little or no modulation with attention. At later times, both spatial and feature-selective attention had robust effects in both ROIs, and these effects were multiplicative rather than additive. In the following analyses we consider how these effects vary across classifications of varying feature difference, and we test for evidence of information exchange between the occipital and frontal ROIs.

### Decoding object features: effect of physical difference between stimuli and task difficulty

Next we considered how classifier performance varied with the physical difference in the stimuli being discriminated (i.e. with task difficulty). Our design included stimuli that were far apart in feature space (e.g. ‘strongly red’ vs ‘strongly green’) and stimuli that were close in feature space (e.g. ‘strongly green’ vs ‘weakly green’). Since we included 4 steps along both color and shape dimensions, the pairs of object stimuli that classifiers were trained to discriminate could be either 1, 2 or 3 steps apart along either dimension. These pairs also differ in task difficulty: for those that are 3 steps apart the stimuli being discriminated were both from ‘easy’ trials, while those of 1 or 2 steps difference contained at least one stimulus from a ‘hard’ trial. In Figure 4 we separately consider classifier performance for pairs of different step size separation, where participants were attending to the stimulus feature and location (pairs of different step size were averaged in Figure 3).

**Figure 4:**
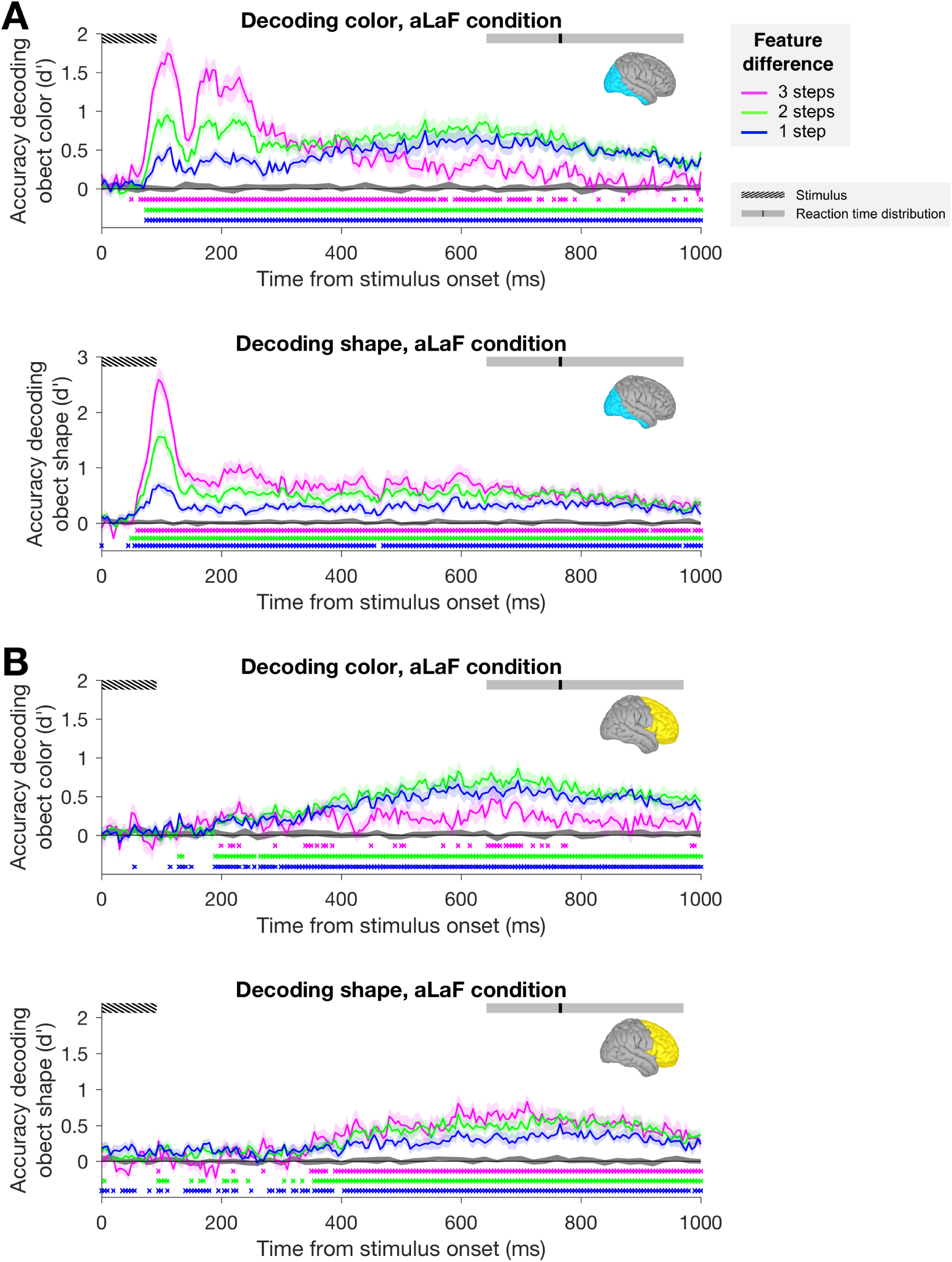
Effect of feature step size on the decoding of object color (upper) and shape (lower) for the occipital (A) and frontal (B) ROIs. In both cases, classifiers were trained to discriminate the shape or color of the object at the attended location, when participants were performing the task relevant to the decoded feature (aLaF condition). Shaded gray region around x-axis indicates the 95% confidence intervals of the same classifications when performed on permuted data (chance performance level, averaged across classifications). Colored crosses below the plot show time points at which classifier accuracy was significantly above the chance performance level (*p* < 0.05 in a one-tailed *t*-test of the between-subject mean, FDR corrected at *q* < 0.05 for multiple comparisons across time points). Shaded error bars indicate the 95% confidence interval of the between-subject mean.

For both decoding of object color and shape, in the occipital ROI performance at the early peak (at around 100 ms after stimulus onset) clearly increased with increasing step sizes. This is consistent with the classifier performance at this early time being driven by predominantly stimulus-driven neural responses in these cortical visual areas. In the case of decoding object shape this ordering persisted throughout the first 1000 ms after stimulus onset for the occipital ROI, and was also seen in the frontal ROI when classification performance emerged.

For decoding object color in the occipital ROI, the order continued until around 350 ms after stimulus onset, when classifier performance on the ‘strongly red’ vs ‘strongly green’ discrimination decreased while classifier performance on 1 or 2 step discriminations increased. Similarly, when classification of color emerged in the frontal ROI performance was weakest for stimuli that were 3 steps apart. The weaker classifier performance at later time points for ‘strongly red’ vs ‘strongly green’ could be related to the participants taking less time to decide their response when judging color on easy trials compared with hard trials. However, this explanation does not account for why there was not a similar effect for decoding of object shape, where reaction times and accuracy on the easy and hard tasks were comparable to that for color. Another possibility is that for the ‘easy’ color trials the participants’ decision was based on neural signals related to the categorization of object color, by an area such as VO (Mullen et al., 2007) or a more anterior area along the ventral temporal processing stream (Lafer-Sousa et al., 2016), with little involvement of frontal areas. Whereas for the more difficult color task trials, and for the shape task, which is unlikely to correspond to a feature dimension of relevance in the occipital cortex, there could have been more involvement by prefrontal areas, which would be consistent with the higher classifier performance in the frontal ROI in these cases.

The relationship between step size and classifier performance was remarkably consistent across the occipital and frontal ROIs. Classifier performance in the frontal ROI did not include the early peak, suggesting that there was good separation of the signals from these different brain regions. But when classifier performance emerged in the frontal ROI the occipital and frontal ROIs showed a very similar pattern of variation across step size, consistent with functional connectivity between these ROIs and the ongoing transfer of stimulus-related information between these brain regions.

### Frontal influence on the occipital representation of object shape and color

To characterize the exchange of stimulus-related information between the occipital and frontal ROIs we used an information flow analysis (Goddard et al., 2016). Since we have fine temporal resolution measures of each pairwise classification, in each attention condition, we used the pattern of classification performance across these measures as a summary of the structure of representational space at each timepoint, and tested for evidence of Granger causal interactions between the ROIs (see Methods for details). Note that by applying this analysis to patterns of classification accuracy (unlike typical Granger causality analyses, which are applied to raw signals), we are not simply testing for evidence of connectivity between brain regions, but are specifically testing for evidence of the exchange of stimulus-related information between areas.

The results of this analysis are plotted in Figure 5. For both color and shape, we found that the earliest time points were dominated by feedforward information flow (FF*>*FB), consistent with the early visual responses in occipital cortex being relayed to frontal regions. These early periods where feedforward information flow dominated were followed by periods of feedback information flow, starting at 285*ms* and 185*ms* for color and shape respectively. In both cases, the information flow is biased towards the feedback direction until ∼400*ms* after stimulus onset. Interestingly, for both color and shape the timing of the feedback information flows align with the onsets of the largest differences in stimulus decoding across attention condition, despite the later onset of these effect for color than for shape. This is seen in Figure 5**B**, where the large divergence between the dark red line (aLaF condition) and the other conditions starts around the onset of the first red region (FB*>*FF), for both color (upper panel) and shape (lower panel). This is compatible with the suggestion that frontal feedback to occipital regions drives the larger attentional effects observed later in the timecourse.

**Figure 5:**
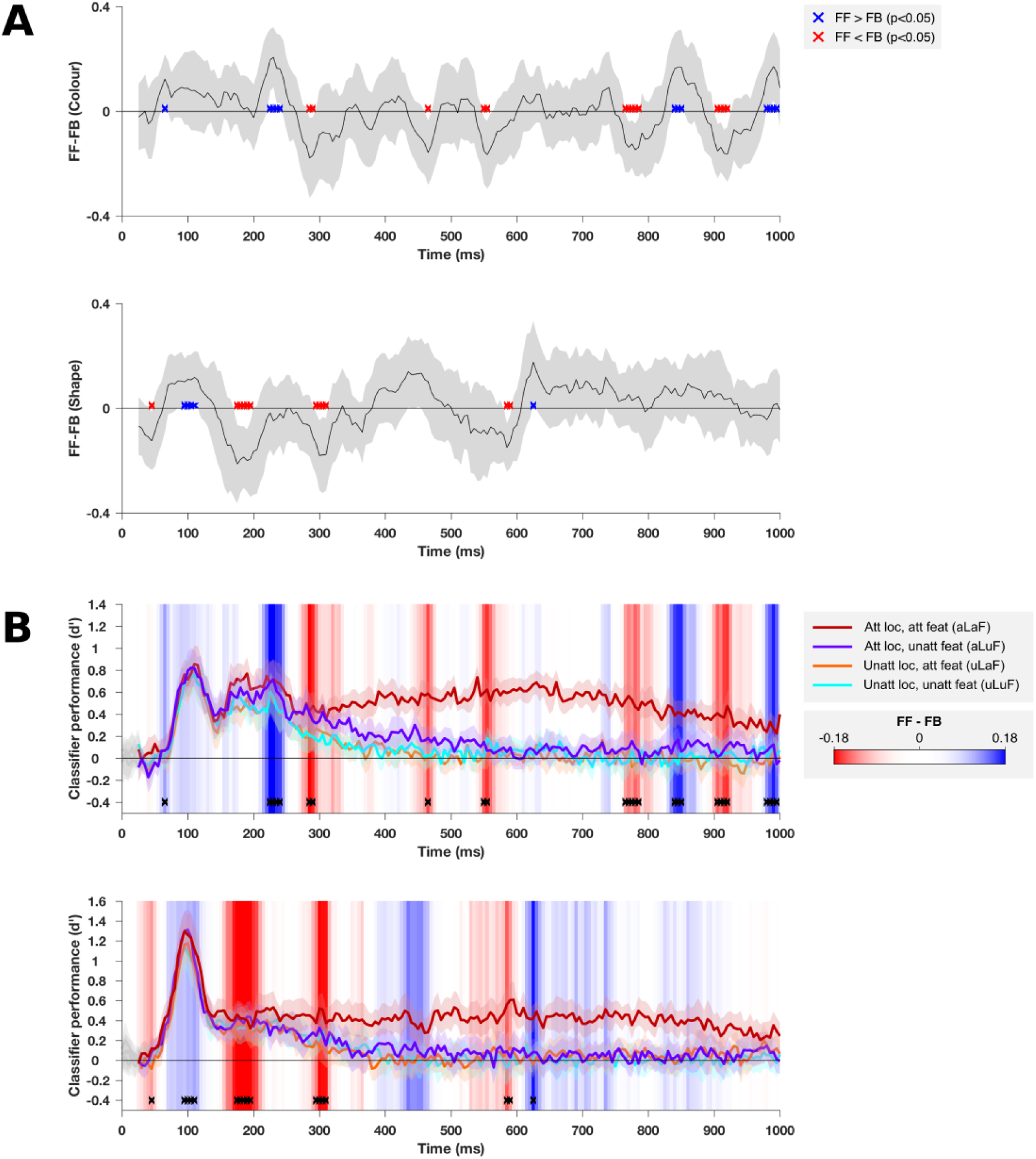
Analysis of feedforward and feedback interactions between occipital and frontal cortices. **A** FF (see Eqn 3) minus FB (see Eqn 4) based on classification performance on decoding stimulus color (upper plot) and shape (lower plot). Time points at which the difference is significantly above or below zero (FF>FB, or FF<FB) are shown in blue and red respectively (*p*-values based on bootstrapped distribution, FDR corrected to *q*<0.05). Shaded error bars indicate the 95% confidence interval of the between-subject mean. In **B** the occipital classification performance in each attention condition is replotted from Figure 3A. The background of the plot is colored according to the data from **A**, as indicated by the colorbar. Time points where FF-FB was significantly different from zero are also replotted, here with black crosses.

The timing differences between color and shape also shed light on the nature of these feedfoward and feedback information flows. For color the early period of FF*>*FB persisted later than for shape (until 240*ms* and 115*ms* after stimulus onset respectively). This extra period of feedforward information flow for color appears to correspond to the second early peak in decoding performance (∼ 165–240*ms* after stimulus onset), and could be related to higher-order processing of color information by occipital cortex at this time, such as the ventral temporal occipital areas (Mullen et al., 2007; Lafer-Sousa et al., 2016). Conversely, since the shape dimension we constructed for this study is highly artificial and unlikely to correspond to a feature dimension of relevance in the occipital cortex, it could be that the earlier feedback signal in this case is related to the frontal cortex’s involvement in storing information about the shape task and in modifying the responses of occipital areas in such a way that the object’s position along the shape dimension can be read out.

Note that while our results are consistent with a late dominance of feedback from frontal to occipital regions, it is possible that the feedback could originate in another area. As with any correlation, it is possible that our partial correlations reflect correlation with another (untested) area. It is also possible that our source reconstruction did not accurately isolate frontal and occipital regions, and that either of these include signals from nearby regions. However, note that if, for example, any parietal signals were present in both frontal and occcipital ROIs, or in the unlikely event that frontal signals were present in the occipital ROI or vice versa, this would tend to reduce the measures of feedfoward and feedback information flows, rather than introduce false positives, making this a conservative analysis. Indeed, the presence of significant feedfoward and feedback information flows provides evidence that the ROIs were well segregated from one another, as does the absence of early classification performance in the frontal ROI.

Later oscillations between feedforward and feedback information flows (>400*ms* after stimulus onset) are more difficult to interpret. Before the median response time (690–820*ms* across conditions) there is a period with a trend towards feedforward information flow for shape (400–500*ms*), but not for color. This may reflect the ‘read-out’ of object shape from occipital cortex, after the occipital responses have been modified by the earlier feedback from frontal cortex: future work may explore this possibility.

### Differential effects spatial and feature-selective attention across feature step size

Figure 3 shows the effects of both the attended location and attended feature on the decoding of object features, and Figure 4 shows that decoding accuracy also varied with how far apart the stimuli were along the relevant feature dimension. We next asked whether there was an interaction between these effects. We reasoned that if feature-selective attention sharpens the population response to the attended feature while spatial attention does not, then they would likely produce qualitatively different patterns of enhancement across stimulus pairs of varying feature difference.

To predict the direction of such an interaction, we used a normalization model of attention (Reynolds and Heeger, 2009) to model the effects of spatial and feature-selective attention on classifier performance. A number of groups have proposed models including normalization to describe the effects of attention on neuronal response properties (Reynolds and Heeger, 2009; Boynton, 2009; Lee and Maunsell, 2009). The normalization model of Reynolds and Heeger 2009 predicts that neuronal responses are given by a stimulus drive that is divided (normalized) by a suppressive drive that varies with the stimulus drive. In the model, the effect of attention on neuronal responses is an ‘attention field’ that varies with spatial position and the stimulus feature dimension, to incorporate the effects of both spatial and feature-based attention. The attention field affects the stimulus drive, and in turn the suppressive drive. Depending on the relative sizes of the stimulus and the attention field, the model can predict changes in both response gain and contrast gain in the response to the attended stimulus. The model also accounts for the sharpening of tuning curves across the visual field with feature-based attention (Martinez-Trujillo and Treue, 2004).

Here we tested whether this normalization model could also predict the effects of spatial and feature-selective attention for our population-level measures of stimulus-related information. Normalization models are based on the average effect of attention on the responses of single neurons, ignoring the heterogeneity of effects across neurons, and the effects of factors such as signal and noise correlations (Sprague et al., 2015; Moreno-Bote et al., 2014). We tested whether this model was useful for predicting patterns of classifier performance despite these simplifications. The model predictions for our experimental design are illustrated in Figure **6A-B**. Figure 6**A** shows the effects of spatial and feature-selective attention on the population response for an example set of parameters, illustrating the predicted sharpening of the population response with feature-selective attention, compared to a more general facilitation across the population response with spatial attention. Details of the model predictions, including further illustrations, are found in the Methods section (see Figure 7). Since the model is descriptive (Reynolds and Heeger, 2009), with a large number of free parameters, we systematically generated model predictions for a wide range of model parameter sets, 172,800 in total. Across these different parameter sets, there was variation in the predicted magnitude of the effects of spatial attention and feature-selective attention, and there was also variation in which stimulus pair feature distances (step sizes) showed the greatest enhancement. However, when compared with spatial attention, feature-selective attention tended to produce relatively more enhancement of small stimulus feature differences than larger ones, as seen in the average difference across all model parameter sets (Figure 6**B**). As seen in Figure S2, a majority of model parameter sets (83%) showed this qualitative pattern of relative enhancement across attention types. Furthermore, there were some combinations of spatial and feature attention excitatory and inhibitory widths for which this same qualitative pattern was found for all 400 combinations of the remaining model parameters (bright red cells in Figure S2).

**Figure 6:**
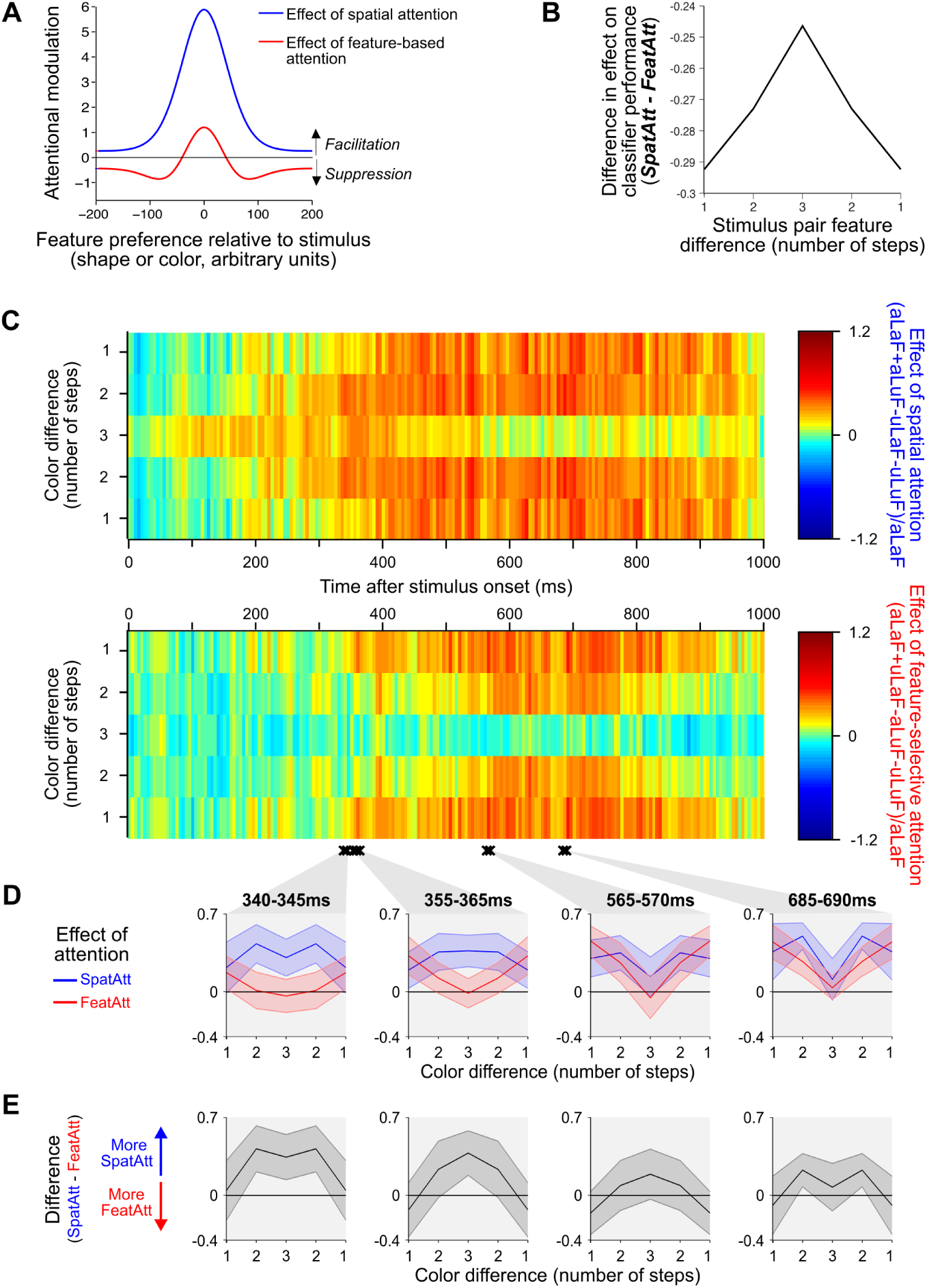
Effects of spatial and feature-selective attention on the decoding of object color in the occipital ROI. **A**: The predicted effects of spatial and feature-based attention on a population of neuronal responses, for an example set of model parameters. According to the model, spatial attention tends to boost the response of all neurons as a multiplicative scaling of the original response, while feature-based attention produces both facilitation of neurons which prefer the attended value, and suppression of neurons preferring nearby values, which leads to sharpening of the population response around the attended value. **B**: Predicted difference between the effects of spatial (**SpatAtt**, Eqn 1) and feature-selective attention (**FeatAtt**, Eqn 2) on classifier performance across pairs of stimuli with different feature differences, averaged over all 172,800 sets of model parameters we tested. **C**: The effects of spatial attention (upper plot) and feature-selective attention (lower plot) on decoding of stimulus color were calculated by taking the difference in classifier accuracy (d’) between the relevant attended and unattended conditions, normalized by the accuracy in the aLaF condition at each time point, for each step size (see Equations 1 and 2). Data from three epochs of interest were averaged and plotted in the insets below (**D**). In **E** the difference between the two attention effects (from the same time points as in **E**) are plotted, and p-values indicate the result of the significance of the interaction between attention type and step-size in each case. The difference values plotted in **C** correspond to the prediction from the model in **B**. Shaded error bars indicate the 95% confidence interval of the between-subject mean.

If feature-selective attention especially enhances the discrimination of small differences along that feature dimension, then we should see a larger effect of feature-selective attention (compared with spatial attention) for pairs of stimuli that differ by only one step, rather than 2 or 3, along the relevant feature dimension. That is, we should see the qualitative pattern from Figure 6**B** in our data. Alternatively, if spatial and feature-selective attention produce qualitatively similar enhancements in the population representation of the stimulus features, we would expect this difference measure (**Diff = SpatAtt-FeatAtt**) to be constant across stimulus step size.

To test this prediction, for stimulus pairs of each step size difference we calculated metrics summarizing the effects of spatial attention (**SpatAtt**, Eqn 1 in Methods) and feature-selective attention (**FeatAtt**, Eqn 2), as shown in Figure 6**C**, for the decoding of color in the occipital ROI. In Figure 6**C-E** and in subsequent figures we plotted data as ‘tuning curves’ across step size, mirror-reversing the data from 1 and 2 steps difference to visually highlight differences between spatial and feature-selective attention in their influence on the shape of these curves. For all statistical analyzes we used data without the mirror reversals.

While our key prediction concerns the difference between **SpatAtt** and **FeatAtt**, in order to give a more complete depiction of the data we plotted these two metrics separately in Figure 6**C**, including data from every step size and time point. In these color plots, cyan to lime indicates that there was little or no effect of attention on classifier performance, while yellow through to red indicates a small to large increase in discriminability. While it is possible for the metrics to have a negative (dark blue) value, which would indicate decreased classifier performance with attention, this was not seen in the data.

If spatial and feature-selective attention produced qualitatively similar effects on neural responses, then the plots in Figure 6**C** should look similar, and the regions of yellow-red should have a similar shape. Instead, visual comparison of the plots in Figure 6**C** reveals differences between the two types of attention in their effects on decoding of color in the occipital ROI. Consistent with the data in Figure 3, the effect of spatial attention emerges earlier than that of feature-selective attention: at « 200 ms there is a band of yellow for spatial but not feature-selective attention. Critically, there was also a systematic difference between spatial and feature-selective attention in their relative effects on classifier performance across step size. In 6**C** this is seen most clearly in the ‘convex’ versus ‘concave’ shape of the yellow-red regions from 300 ms after stimulus onset in the upper and lower plots. Furthermore, while spatial attention tended to produce the greatest increase in classifier performance (the largest red area) for stimuli separated by 2 steps in feature space, feature-selective attention tended to produce greatest enhancement for stimuli separated by only 1 step along the relevant feature dimension (the stimulus pairs that were most similar).

To identify times at which spatial and feature-selective attention differed in their effects across step size we performed a 2-way ANOVA, with subject as a random factor, at each time point. Clusters of time points at which there was a significant interaction between attention type and step size (*p* < 0.05, at least 2 consecutive time points) are indicated by the black crosses in Figure 6**C**. The earliest cluster began after the second peak in classification performance, at 340ms after stimulus onset. To visualize the interaction at these times, and in order to plot the inter-subject variability, for each cluster we plotted the average effect of spatial and feature-based attention (Figure 6**D**), including 95% confidence intervals of the between-subject mean. For every cluster of timepoints for which there was a significant interaction between attention type and step size the effect went in the same direction: spatial attention had a greater effect than feature-selective attention at the largest step size, while feature-selective attention had a larger effect than spatial attention at the smallest step size. This is illustrated most clearly in the difference plots (**SpatAtt-FeatAtt**) of Figure 6**E**. As an additional control, we confirmed that the same pattern of results persists when excluding participants with any bias toward the attended location in their average fixation location (Figure S4). These data suggest a robust qualitative difference between spatial and feature-selective attention in the way they enhance the color information in occipital areas.

For the decoding of shape in the occipital ROI, the effects of spatial and feature-selective attention were more uniform across step sizes (see Figure S5), and there were no clusters of time points with a significant interaction between attention type and step size. This was also true for the frontal ROI, for decoding of both color and shape (data not shown). In order to test if there any interaction between attention types and step size for object shape when data from the entire brain was included, we also calculated **SpatAtt** and **FeatAtt** for the decoding of object shape based on sensor data (before any source localization). In this case, there were 2 clusters of consecutive time points where there was a significant interaction between attention type and step size (Figure S6), and the earliest of these began at 365ms after stimulus onset. Notably, where these interactions occurred, the effects were also in the predicted direction, despite variation in the effect of step size on decoding color and shape (Figure 4). This suggests a general qualitative difference between spatial and feature-selective attention in the way they enhance the information that is carried by neural population codes, which aligns with that predicted by a normalization model.

## Discussion

Attentional selection is critical for fast and accurate processing of behaviorally relevant visual information. There are different methods by which we can select a subset of visual information for further processing, but the extent to which these are implemented by similar or different neural processes, and how these attentional effects interact, remains unclear. Spatial and feature-selective attention have rarely been directly compared within the same experiment, and to our knowledge this is the first test of two key predictions regarding their interaction: that spatial and feature-selective attention interact in a multiplicative way in their effects on neural coding, and that they induce qualitatively different patterns of enhancement across fine and coarse feature differences. We found that a normalization model of attention, designed primarily to account for the effects of attention on individual neurons, predicts these effects of attention on the information carried by a population neural signal.

Previous neuroimaging work has revealed some of the effects of spatial (Brefczynski and DeYoe, 1999; Jehee et al., 2011; Guggenmos et al., 2015; Sprague and Serences, 2013) and feature-selective (Corbetta et al., 1990; Chawla et al., 1999; Saenz et al., 2002; Serences and Boynton, 2007; Saproo and Serences, 2014; Jackson et al., 2016; Vaziri-Pashkam and Xu, 2017) attention at a population level. Like some previous fMRI studies, we used classifier accuracy as an intuitive means of measuring the effects of attention: using classifier accuracy as a proxy for the amount of information that is potentially available in the neural response. Here we applied this decoding approach to MEG data, which allowed us to explore the timecourse of these effects using the millisecond resolution of MEG. We found evidence that both spatial and feature-selective attention boost the stimulus-related information in the population response, and we were able to measure these effects in both frontal and occipital regions. In both frontal and occipital regions, the effects of spatial attention emerged earlier than those of feature-selective attention. Through our information flow analysis of Granger-causal relationships between occipital and frontal regions, we found that stimulus-related activity in frontal regions influenced occipital representations from as early as 185*ms* after stimulus onset, and that the onset of this influence coincided with the largest magnitude attentional effects in occipital regions. In addition, we found evidence confirming two predictions relating to how the effects of spatial and feature-based attention interact, and how they differ in their relative enhancement of the discriminability small versus large stimulus feature differences. We consider each of these findings below.

### Earliest responses in occipital areas modulated by spatial but not feature-selective attention

For the decoding of both color and shape, we found that spatial attention had only a small effect and feature-selective attention had no significant effect on the initial peak of classifier performance in the occipital ROI (∼100*ms*), but much larger effects at later times. The effect of feature-selective attention on occipital stimulus representation was only significant from 335–385*ms*: at least 200*ms* after the effect of spatial attention. Furthermore, while there was a small effect of spatial attention around the initial peak in classifier performance (∼100*ms* after stimulus onset) there was no significant effect of feature-selective attention, consistent with another report that the earliest occipital responses are not affected by feature-based attention (Bartsch et al., 2017). This finding that spatial attention effects preceded those of feature attention is consistent with previous results from electrophysiological recordings in V4 and FEF (Zhou and Desimone, 2011; Bichot et al., 2015), although the delay observed here is longer than in this previous work. For both features, feature-selective attention had an impact on classifier performance in the occipital only after feedback from the frontal ROI began to dominate the information flow (FB*>*FF). Since information flow analysis specifically measures the exchange of stimulus-related information, this result suggests that the effects of feature-selective attention in occipital cortex may rely on feedback of stimulus-related information from frontal areas.

The degree to which subjects are engaging attention prior to stimulus onset could also have contributed to the pre-stimulus decoding of attentional task for spatial but not feature-selective attention (Figure 2) and to the earlier effects of spatial attention, relative to feature-selective attention on the stimulus representation (Figure 3). For example, it may be easier to prepare to attend to a location than to prepare to attend to a feature dimension. A previous study reported that feature-based attention can modulate event-related potentials (ERPs) much earlier than in our data, within 100*ms* of the stimulus onset (Zhang and Luck, 2009) (in contrast to 335 – 380*ms* onset in the our results). This discrepancy may reflect a difference between feature-based attention (attending to a feature value, e.g. ‘red’) and feature-selective attention (attending to a feature dimension, such as ‘color’). Another critical difference between these studies is in stimulus design: Zhang and Luck (2009) recorded responses to a flashed probe stimulus of red or green dots while subjects attended to dots of one color in another covertly attended stimulus, where dots of both colors were always present. In our experiment, the stimuli were always preceded by a blank screen, so that subjects were planning to attend to a particular stimulus feature rather than already attending to it. In our data, decoding of attention condition became much stronger once stimuli appeared and the participants were actively performing the task. We hypothesize that these stimulus differences account for the later onset of feature-selective attention’s effect on stimulus representation here, and that the early effects of feature-based attention reported by Zhang and Luck (2009) are only present when the subject is already engaged in attending to one feature value (or suppressing the irrelevant feature value, see Moher et al. 2014; Andersen and Müller 2010).

### Information flow analysis: the role of frontal feedback in attentional modulation

The earliest responses of the occipital cortex showed little modulation with attentional condition, consistent with a stimulus-driven response. Shortly after these initial responses there were large effects of both attention types: attention changes the stimulus information representated by the population response in occipital cortex. What regions drive the effects of attention on the occipital population response? Within occipital cortex, previous work suggests that attentional effects are present first in higher-order visual areas that induce a top-down modulation of earlier areas (Buffalo et al., 2010), but this leaves open the possibility that effects in higher-order visual areas are driven by another region. Our information flow analysis suggests a contribution from frontal areas, with stimulus-related information coding in occipital cortex appearing to follow from the information coding in the frontal lobe shortly beforehand. A class of models of prefrontal function converge on the proposal that prefrontal cortex implements cognitive control by affecting processing in more specialised cortices (Duncan, 2001; Desimone and Duncan, 1995; Dehaene et al., 1998; Miller and Cohen, 2001). By tracking the dynamics of information exchange between frontal and occipital cortex we were able to test this suggestion and resolve the timecourse of the proposed top-down effects.

We found that information flow was initially dominated by feedforward propagation of information from occipital to frontal lobe, then later dominated by information flowing in the opposite direction, with information coding in the frontal ROI predicting subsequent information coding in occipital cortex (see also, Goddard et al. 2016; Karimi-Rouzbahani 2018). Moreover, the onset of feedback dominating the flow of information between frontal and occipital cortex corresponded to the time at which the occipital lobes showed a divergence between task-relevant and task-irrelevant information. For decoding color, where there was a second early peak in classifier performance, this period was later (285*ms*) than for decoding shape (185*ms*), but in both cases it aligned with the time at which information processing in the occipital lobes became dominated by the task-relevant information (classifier performance in the attended location, attended feature condition remained steady or increased, while performance in other conditions was strongly attenuated).

Our finding that prefrontal cortex appears to shape responses in occipital areas is consistent with work demonstrating that the responses of frontoparietal regions contain stimulus-related information (for example, Freedman et al. 2001), that increases with spatial (Woolgar et al., 2015) and feature-selective (Jackson et al., 2016) attention, and that attentional effects in frontal cortices precede those in sensory cortex (e.g. Lennert and Martinez-Trujillo 2013). One prominent model of prefrontal cortex function (biased competition model Desimone and Duncan 1995; Duncan 2006) proposes that the prefrontal cortex biases processing in more specialized (visual) cortices in favor of task-relevant information. In line with such a proposal, our data suggest that after an initial feedforward sweep of information, feedback from frontal to occipital cortices drives the selective representation of information in the occipital cortex.

Future work could build on these findings in two ways. First, we chose not to resolve into more fine-grained parcellations of the frontal lobe here because of the limitations of not having individual MRI scans and concerns about the inverse problem. This presents and interesting avenue for future work using the methods described here, perhaps using concurrent EEG and individualized MRI scans to constrain the inverse problem. Second, with better source estimation it would be interesting to examine the role of other brain regions, particularly the parietal lobe (which is known to have important roles in attention, e.g. Duncan 2010; Woolgar et al. 2011; Hebart et al. 2018; Jerde et al. 2012). In the context of information flow analyses such finer parcellations could identify cases in which correlations between two brain regions are likely mediated by both areas correlating with a third.

### Differential effects of spatial and feature-selective attention as predicted by a normalization model of attention

Much of our knowledge of spatial and feature-selective attention comes from studies that have investigated their effects in separate experiments. As such, the results presented here provide valuable new insight into how these two types of attention interact. We found that where there were effects of both types of attention there also tended to be an interaction between them, which is consistent with a multiplicative rather than an additive combination of attentional effects. In the normalization model of attention presented by Reynolds and Heeger (2009), they modeled all but one of the results with a multiplicative rather than additive interaction^1^

We used results from single-unit work to predict how differences in the effects of spatial and feature-selective attention might manifest in population-level codes for stimulus features. Specifically, we predicted that feature-selective attention would produce relatively more enhancement of classifier performance for small feature differences than for large feature differences, as compared with the effects of spatial attention. We confirmed this intuition by using a normalization model (Reynolds and Heeger, 2009) to generate predictions for our data. Normalization models of attention are primarily based upon the electrophysiological study of the effects of spatial and feature-based attention on tuning of individual cells, yet here we demonstrate that the same model can account for population level data, and can be extended to predict the effects of feature-selective attention. It is particularly important that we understand how the effects of attention manifest at a population level since there are significant effects at a population level that cannot be captured by measuring the tuning curves of individual cells (Sprague et al., 2015; Cohen and Maunsell, 2009). The results of our classification analyses based on the MEG data revealed that spatial and feature-selective attention have distinct effects on stimulus-related information coding at a population level, and these differences were consistent with the predictions of the normalization model.

The fact that classifier performance was consistent with the predictions of the normalization model does not definitively identify what information the classifier analysis is using to decode stimulus color and shape, which is difficult to pin down in any case where classifiers are used to measure stimulus-related information from neuroimaging data (Carlson et al., 2018). However, this result suggests that the information that is accessible to the classifier varies in signal strength in a manner that is consistent with what we expect based on the effects on single-unit tuning predicted by the normalization model. Additionally, differences between decoding of color and shape are broadly consistent with color (but not ‘X-shaped-ness’) being a feature dimension that is explicitly encoded by visual cortex. We found the most marked difference between the attention types in the decodability of stimulus color in the occipital ROI. Of the two feature dimensions we manipulated (shape and color) it is more plausible for color that there are single-units with response functions that approximate those included in the normalization model. Neurons in a range of visual cortical areas are tuned for color (for example, Komatsu et al. 1992; Hanazawa et al. 2000), and attention to color is a form of feature-based and feature-selective attention that has been investigated in single-unit work (for example, Motter 1994; Bichot et al. 2005; Chen et al. 2012). In contrast, the shape dimension (from ‘X-shaped’ to ‘non-X-shaped’) is an artificial, more complex dimension than color. It is possible that this dimension could align with the feature selectivity of some neurons in an area with intermediate to high level shape selectivity, such as the in area V4 (see review by Pasupathy 2006), but it is unlikely that there is population tuning for this shape dimension in the same way that we expect a population code for the color dimension. Although the tuning differences between spatial and feature-selective attention were weaker for shape than for color, where these differences were significant (in the sensor-level decoding) the effect was in the same direction as for color. This suggests that a population tuning curve framework may be helpful for understanding the effects of attention on arbitrary, higher level feature dimensions as well as for lower-level ones.

Normalization models of attention can account for a range of the effects of attention observed at the level of a single neuron (Boynton, 2005; Reynolds and Heeger, 2009; Boynton, 2009; Lee and Maunsell, 2009). Although designed to model single-neuron effects, these models can be used to predict attention-based changes in the information carried by the population response, such as in the implementation used in the present study. For both single-unit and population responses these models are primarily descriptive rather than quantitative, but in selecting ranges of model parameters we considered parameters that are feasible for single-unit responses and found that these same parameters could account for population-level effects. Our results demonstrate that the same principles that describe phenomena at the single-unit level, such as multiplicative scaling in spatial attention, and sharpening of the population response in feature-selective attention, can account for population level changes, particular in the encoding of color by occipital areas. Notably, the normalization model successfully predicted these population-level effects despite the fact that the model does not incorporate any heterogeneity of effects across neurons, nor any effects of signal or noise correlations, which could have caused differences between single-unit and population-level effects (Sprague et al., 2015; Moreno-Bote et al., 2014). This opens the possibility of using such models as an explanatory bridge between levels of description: if future work constrains model parameters for the normalization model at either the single-unit or the population level this may generate predictions that can be tested at other, to further characterize the similarities and differences between these levels of description. When model parameters are further constrained by data, another direction for future work is to test quantitative as well as qualitative predictions of these models.

## Conclusions

We used multivariate pattern analysis of MEG recordings to measure the effects of spatial and feature-selective attention on the amount of stimulus-related information decodable from large populations of neurons. We manipulated both spatial and feature-selective attention simultaneously in order to compare these attention types within the same dataset, and to test how these attention types interact in their effects on population-level representation of visual stimuli. We found that both spatial and feature-selective attention enhanced the representation of visual information and that the two types of attention interacted in a multiplicative way to yield an adaptive neural representation which prioritised the task relevant feature of the attended object. An information flow analysis suggested that the largest attentional effects in occipital areas may be driven by feedback from frontal areas.

We further found that modelling the distinct effects of spatial and feature attention at the level of single cells predicted the qualitative differences between spatial and feature-selective attention in our population level recordings. The success of the modelling was remarkable given that the model only included the effects of attention on tuning properties, without modelling, for example, any influence of attention on the correlation structure of the population. Specifically, consistent with a normalization model of attention in which feature-selective attention results in tuning curve sharpening and spatial attention predominantly yields response gain, we found that for decoding of color in occipital cortex, feature-selective attention produced more enhancement of the neural representation of small stimulus feature differences than spatial attention did, while spatial attention resulted in greater discrimination of large stimulus feature differences.

Our ability to direct our attention to different locations and to different features of the environment appears to rely on interacting attentional mechanisms that induce qualitatively distinct changes in population-level neural responses in sensory cortices.

## Materials and Methods

### Participants

20 volunteers (14 female, 6 male) participated in this study, and were paid $50 as compensation for their time. Participants ages ranged from 18-32 years (mean 22.4 years). All were right-handed, had normal or corrected to normal vision, had no history of neurological or psychiatric disorder, and were näıve to the purposes of the study. All participant recruitment and experiments were conducted with the approval of the Macquarie University Human Research Ethics Committee.

### Visual stimuli

Visual stimuli were generated and presented using Matlab (version R2014b) and routines from Psychtoolbox Brainard (1997); Pelli (1997). We created novel object stimuli that varied in color and in their shape statistics (see Figure 1**B**), using custom code. The shapes were variants of ‘spikie’ stimuli used in previous work (Op de Beeck et al., 2006; Woolgar et al., 2015; Jackson et al., 2016). All our ‘spikie’ shapes had a common almost spherical body and 16 spikes varying in location, length and orientation. All shapes were rendered with diffuse illumination and a direct (upper left) illuminant source, and presented on a black background. We varied the spike orientation statistics to create four classes of ‘spikie’ objects: strongly ‘X-shaped’, weakly ‘X-shaped’, weakly ‘non-X-shaped’, and strongly ‘non-X-shaped’ (Figure 1**B**). When performing the shape-based task participants categorized the target object as either ‘X-shaped’ or ‘non-X-shaped’. We created 100 unique versions of each shape class by adding random variation in the spike locations, lengths and orientations, to ensure that participants could not perform the task by attending to a single feature, and to encourage them to attend to the object’s overall shape.

In color, there were also four classes: ‘strongly red’, ‘weakly red’, ‘weakly green’ and ‘strongly green’ (Figure 1**B**). When performing the color-based task participants categorized the target object as either ‘reddish’ or ‘greenish. Each object had a maximum luminance of 108.1 *cd*/*m*^2^, and constant u’v’ and xy chromaticity coordinates (Wyszecki and Stiles, 1982). The chromaticity coordinates were as follows; strongly red u’v’: 0.35, 0.53 (xy: 0.56, 0.38); weakly red u’v’: 0.27, 0.54 (xy: 0.50, 0.44); weakly green u’v’: 0.23, 0.55 (xy: 0.45, 0.48) and strongly green u’v’: 0.16, 0.56 (xy: 0.36 0.57). The weak red and weak green colors were defined as lying on a line joining the strong red and strong green coordinates in u’v’ space, and their distance from the line’s midpoint was 30% of the distance between the midpoint and the relevant endpoint.

During MEG sessions, stimuli were projected through a customized window by an InFocus IN5108 LCD back-projection system (InFocus, Portland, Oregon, USA) located outside the Faraday shield, onto a screen located above the participant. Participants, lying supine, viewed the screen from 113cm. Individual ‘spikie’ objects each had a central body of 195 pixels (5.8 degrees visual angle [dva]) wide x 175 pixels (5.2 dva) high. Their total size varied with their spikes, but the spikes never reached the border of the object image (403×403 pixels). On each trial, the stimulus included 2 ‘spikie’ object images side-by-side (total size 806 pixels wide x 403 pixels high: 24 x 12 dva). A white fixation cross, with height and width of 1 dva, was drawn in the center of the screen (Figure 1**A**). The display system was characterized in situ using a Konica Minolta CS-100A spectrophotometer and calibrated as described previously (Goddard et al., 2010).

### Experimental design: MEG and eye tracking

MEG data were collected with a whole-head MEG system (Model PQ1160R-N2, KIT, Kanazawa, Japan) consisting of 160 coaxial first-order gradiometers with a 50 mm baseline (Kado et al., 1999; Uehara et al., 2003). Prior to MEG measurements, five marker coils were placed on the participant’s head. Marker positions, nasion, left and right pre-auricular points, and the participant’s head shape were recorded with a pen digitizer (Polhemus Fastrack, Colchester, VT), using a minimum of 2000 points.

Each participant’s MEG data were collected in a single session of approximately 90 minutes, at a sampling frequency of 1000Hz. On each trial participants responded using a Fiber Optic Response Pad (fORP, Current Designs, Philadelphia, PA, USA).

We tracked participant’s eye movements using an EyeLink 1000 MEG-compatible remote eye-tracking system (SR Research, 500Hz monocular sampling rate). Before scanning we tested participants for their dominant eye (usually right), and focused the eye-tracker on this eye.

### Experimental design: participant’s task

Each participant’s MEG session was divided into 8 blocks, where the location of the attended object (left or right of fixation) and the task (reporting the attended object’s shape or color category) was constant within each block. Figure 1**A** illustrates the four different attention conditions. Before the experiment, each participant was familiarized with the ‘X-shaped’ and ‘non-X-shaped’ object categories and completed a training session on a laptop outside the MEG scanner where they practiced the color and shape tasks.

On every trial we presented two objects, one each on the left and right of fixation. We presented the objects simultaneously since both spatial attention (Reynolds and Desimone, 1999; Sundberg et al., 2009) and feature-selective attention (Saenz et al., 2003) effects are stronger when attended and unattended stimuli simultaneously compete for access to perceptual processing. Within each block every pairing of the 16 objects in Figure 1**B** was included once, giving 256 (16×16) trials. These 256 trials were presented in a counterbalanced order within each block, so that objects of each shape and color were equally likely to precede objects of all shapes and colors. A different counterbalanced order was used for each block, and to this sequence of 256 trials the last trial was added to the beginning, and the first trial was added to the end, giving a total of 258 trials in each block. Data from these first and last trials were discarded.

The participant’s task alternated between shape and color on every block, and the location of the attended object alternated after the 2nd, 4th and 6th blocks. Starting location and task were counterbalanced across participants. Within each pair of blocks where the attention condition was the same (e.g. blocks 1 and 5), the buttons corresponding to the two response options were switched, so that response mappings were counterbalanced across blocks.

Every block commenced with an instruction regarding the attended object, the task, and the response mapping for that block. Before the first trial participants were required to identify the response buttons correctly with a keypress. Participants also repeated the eye-tracker 5-point calibration, before the block commenced.

Every trial began with the fixation marker alone while the participant’s fixation was verified using the eye tracker. Participants had to fixate within 1 dva of the fixation marker for at least 300 ms before the stimulus would appear. During the stimulus (maximum 150ms) a 50×50 pixel white square appeared in the bottom right corner of the projected image (outside the stimulus region), which was aligned with a photodetector, attached to the mirror, whose signal was recorded with the MEG signal from the gradiometers. We used the photodiode signal to accurately align MEG recordings with stimulus timing during data analysis. When eye-tracking showed participants were no longer fixating during the 150ms stimulus presentation, the stimulus was removed from the screen. Due to eye tracker variability (e.g. eye tracker missing frames), this resulted in an unexpectedly high number of shorter trials: the median stimulus duration was 92ms, and the first and third quartiles were 64 and 126ms. Since this affected a majority of trials, we included all trials in our analysis, but ran an extra analysis to check that variability in stimulus duration did not account for our results (see below). After stimulus offset, the fixation marker remained white until participants responded to the appropriate task via a button press. After the participant’s response, but no sooner than 1000 ms from the onset of the stimulus, the fixation marker changed for 200 ms to provide feedback: dimming to gray for ‘correct’, or turning blue for ‘incorrect’. After feedback, there was a variable inter-trial interval (300-800ms), which comprised the fixation check for the subsequent trial. We used a variable inter-trial interval to avoid expectancy effects. Across participants, the median reaction time was 0.77s (shape task: 0.78s; color task: 0.75s); on 77% of trials the reaction time was shorter than 1 s and the feedback onset was 1 s. The first and third quartiles of the distributions of reaction times are shown in Figures 2 and 3.

### MEG data analysis: Source reconstruction

Forward modeling and source reconstruction were performed using Brainstorm (Tadel et al., 2011), which is documented and freely available for download online (http://neuroimage.usc.edu/brainstorm). First, we created a model of each participant’s brain by manually aligning the ICBM152 template brain (Fonov et al., 2011) to their head shape using nasion, pre-auricular points, and head shape data. Once aligned, we applied nonlinear warping to deform the template brain to the participant’s head shape, which provides a superior model to an unwarped canonical template (Henson et al., 2009). We generated a forward model for each model by applying a multiple spheres model (Huang et al., 1999) to the individually warped template brain and their measured head location.

Functional data were preprocessed in Brainstorm with notch filtering (50, 100 and 150Hz), followed by bandpass filtering (0.2-200Hz). Cardiac and eye blink artifacts were removed using signal space projection (SSP): cardiac and eye blinks events were identified using default filters in Brainstorm, manually verified, then used to estimate a small number of basis functions corresponding to these noise components, which were removed from the recordings (Uusitalo and Ilmoniemi, 1997). From these functional data we extracted two epochs for each trial: first, a measure of baseline activity (−100 to −1ms relative to stimulus onset), and secondly the evoked response (0 to 1000ms). We used the baseline measures to estimate the noise covariance for each run, then applied a minimum norm source reconstruction to the evoked data. For each source reconstruction, we used a 15,000 vertex cortical surface (standard for the ICBM152 template, with atlas information). Dipole orientations in the source model were constrained to be normal to the cortical surface, the noise covariance was regularized using the median eigenvalue and all other options were set to their default values. We visually inspected the quality of the source reconstruction: the average trial data included an initial ERP at the occipital pole and subsequent ERPs at sources within the occipital cortex but lateral and anterior to the occipital pole, consistent with extrastriate areas along the ventral visual pathway (see Supplementary Figure S1).

### MEG data analysis: Preprocessing and dataset definitions

For classification analyses we generated three datasets: the first included preprocessed data from all sensors, without source reconstruction. The second included sources in occipital, occipito-temporal, and inferior-temporal cortices (‘Occipital’ ROI, 3302 vertices) in the atlas for the ICBM152 template, and the third included frontal and prefrontal cortices (‘Frontal’ ROI, 3733 vertices), as shown in Figure 2**A**.

For each dataset, we extracted data from −100 ms to +2000 ms relative to the stimulus onset of each trial. We then reduced each data set, comprising 2100 ms of data for each of 2048 trials and up to 160 sensors or up to 3733 sources, using PCA. We retained data from the first *n* components which accounted for 99.99% of variance (mean, std *n*: 85.3, 6.9 for frontal ROI; 76.6, 5.8 for occipital ROI; and 157.2, 1.1 for whole brain sensor data) and down-sampled to 200Hz using the Matlab ‘decimate’ function.

### MEG data analysis: Classifier analyses

We used classification analyses to measure the extent to which brain activity could predict attention condition and the color and shape of the stimuli on each trial. For every classification we repeated the analysis at each time point (each 5ms bin) to capture how the information carried by the neural response changed over time: we trained classifiers to discriminate between two categories of trial and tested on held-out data. We report results obtained with a linear support vector machine (SVM) classifier, using the Matlab function *fitcsvm* with ‘KernelFunction’ set to ‘linear’. We also repeated our analyses with a linear discriminant analysis (LDA), using the Matlab function *classify* with ‘type’ of ‘diagLinear’ and obtained very similar results (not shown).

For each classification we created ‘pseudo-trials’ by averaging across trials with the same value on the dimension-of-interest, but with differing values along other dimensions. We used pseudo-trials in order to increase signal-to-noise along the dimension-of-interest (e.g. see Guggenmos et al. 2018; Grootswagers et al. 2017). For example, when classifying the attended location, we took the 4 blocks of 256 trials where the participant attended to the object on the left, and generated 256 pseudo-trials, each the average of 4 trials with one randomly-selected trial from each block. This meant that each pseudo-trial included data from an equal number of trials from the attended feature conditions (attend to color and attend to shape). For each classification we generated 100 sets of pseudo-trials, updating the random assignment of trials for each set, and averaged classification performance across these.

Features that were balanced across pseudo-trials varied with the feature-of-interest being classified. As mentioned above, when classifying attended location pseudo-trials were balanced across attended feature. Similarly, for classifying attended feature pseudo-trials were balanced across attended location. When training classifiers to discriminate object color and shape, we trained and tested within a single attention condition (e.g. attend left, report color), comprising two blocks (512 trials). We trained classifiers separately on each pair of the 4 levels along each feature dimension, at each object location, using pseudo-trials to balance across irrelevant dimensions. For example, when classifying ‘strongly green’ versus ‘weakly green’ objects on the left of fixation, we balanced pseudo-trials across left object shape, and right object color and shape. Since balancing across all 3 of these irrelevant dimensions would not provide sufficient data for classifier training (only 2 pseudo-trials per category), we instead created pseudo-trials that were balanced across 2 of these 3 irrelevant dimensions, and randomized across the third (allowing 8 pseudo-trials per category). As before, we generated 100 sets of the pseudo-trials, each with a different randomization.

Additionally, we repeated this entire process 3 times, balancing across different pairs of irrelevant features. For each of set of pseudo-trials, we trained a classifier using 7 of the 8 pseudo-trials in each condition and tested using the remaining pair of trials, repeating 8 times. We averaged classifier performance across these 8 classification boundaries, and across the 300 sets of pseudo-trials.

For color and shape we performed the classification analysis pairwise for each pair of feature values, then averaged classifier performance across feature differences of the same ‘step size’. Since both dimensions had 4 values, pairs were either 1, 2 or 3 steps apart along the given feature dimension. Pairs 2 or 3 steps apart belonged to opposite categories in the participant’s task (‘greenish’ vs ‘reddish’ and ‘X-shaped’ vs ‘non-X-shaped’). Pairs 1 step apart could be within or across these categories; we did not find any differences between these data (data not shown) so averaged across these when reporting our results.

For all analyzes we expressed average classifier accuracy in d’ (a unit-free measure of sensitivity) which provides an intuitive measure of effect size: a *d′* value of 0 corresponds to no stimulus-related information, which was useful when calculating the effects of spatial and feature-selective attention (below). To test whether classifier performance was above chance performance, we repeated each classification analysis for data where trial labels were randomly permuted. We repeated this 10 times for data from every 4th time bin (one every 20ms). In statistical tests we tested whether the observed classification performance exceeded the average chance performance across time bins. Across classifications, average chance performance varied from d’=0.000 to a maximum of d’=0.015.

Additionally, to predict the effect of variable trial duration, we repeated each classification of stimulus feature using the stimulus state (on or off) at each time point. Across time points, the maximum average classifier accuracy was d’=0.4 for this data, indicating that stimulus variability could have made a small contribution to overall accuracy. However, there was very little difference between this decoding for different attention conditions or across step sizes. When we performed the statistical tests reported in Figures 3 on the trial duration data, the only significant result (effect of attended location for decoding stimulus color) was in the opposite direction (decoding was higher for unattended than attended locations).

For each stimulus classification boundary, we averaged the classifier weights across each set of pseudo-trials to generate an estimate of the classifier weights for each participant’s data, at each time point. The magnitudes of raw classifier weights can vary with both signal strength and noise magnitude, making maps of raw weights difficult to interpret (Haufe et al., 2014). To obtain more informative maps we followed a method used previously (Haufe et al., 2014; Wardle et al., 2016) to transform the classifier weights: For each vector (**W**) of average classifier weights across occipital or frontal vertices, we obtained the transformed weights (**W’**) using the covariance matrix of the *n* pseudo-trials that constituted the classifier training/test data (**cov** (*pseudotrials*), using **W’** = **cov** (*pseudotrials*) * **W**. We averaged these transformed weights (**W’**) across all pairwise comparisons before multiplying the weights by the subject-specific PCA coefficients, and finally averaging across participants.

To summarize the effects of spatial attention (**SpatAtt**) and feature-selective attention (**FeatAtt**), we used the following metrics, based on classifier performance (*d*′) in the attended location, attended feature (*aLaF*) condition, the attended location, unattended feature (*aLuF*) condition, the unattended location, attended feature (*uLaF*) condition, and the unattended location, unattended feature (*uLuF*) condition. For both attention effects, we normalized the effects by the classifier accuracy in the *aLaF* condition to minimize the influence of overall classifier accuracy on the estimates of attention effects.

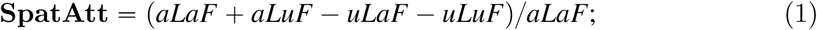

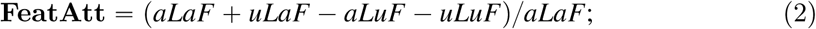

### Modeling the effects of spatial and feature-selective attention on population representations of shape and color

We used a normalization model of the effects of attention at the cellular level to make predictions of how attention would affect stimulus-related information in the population response. Intuitively, we expected that if feature-based attention sharpens the population response to the attended feature, then feature-selective attention should particularly increase classifier performance for stimulus pairs with small feature differences. Conversely, spatial attention, which is not thought to sharpen population responses, should produce relatively more enhancement of classifier performance for larger feature differences. To formalize this intuition we implemented the Reynolds and Heeger (2009) normalization model of attention to generate predictions, as illustrated in Figure 7 and detailed in the Supplementary Methods.

**Figure 7:**
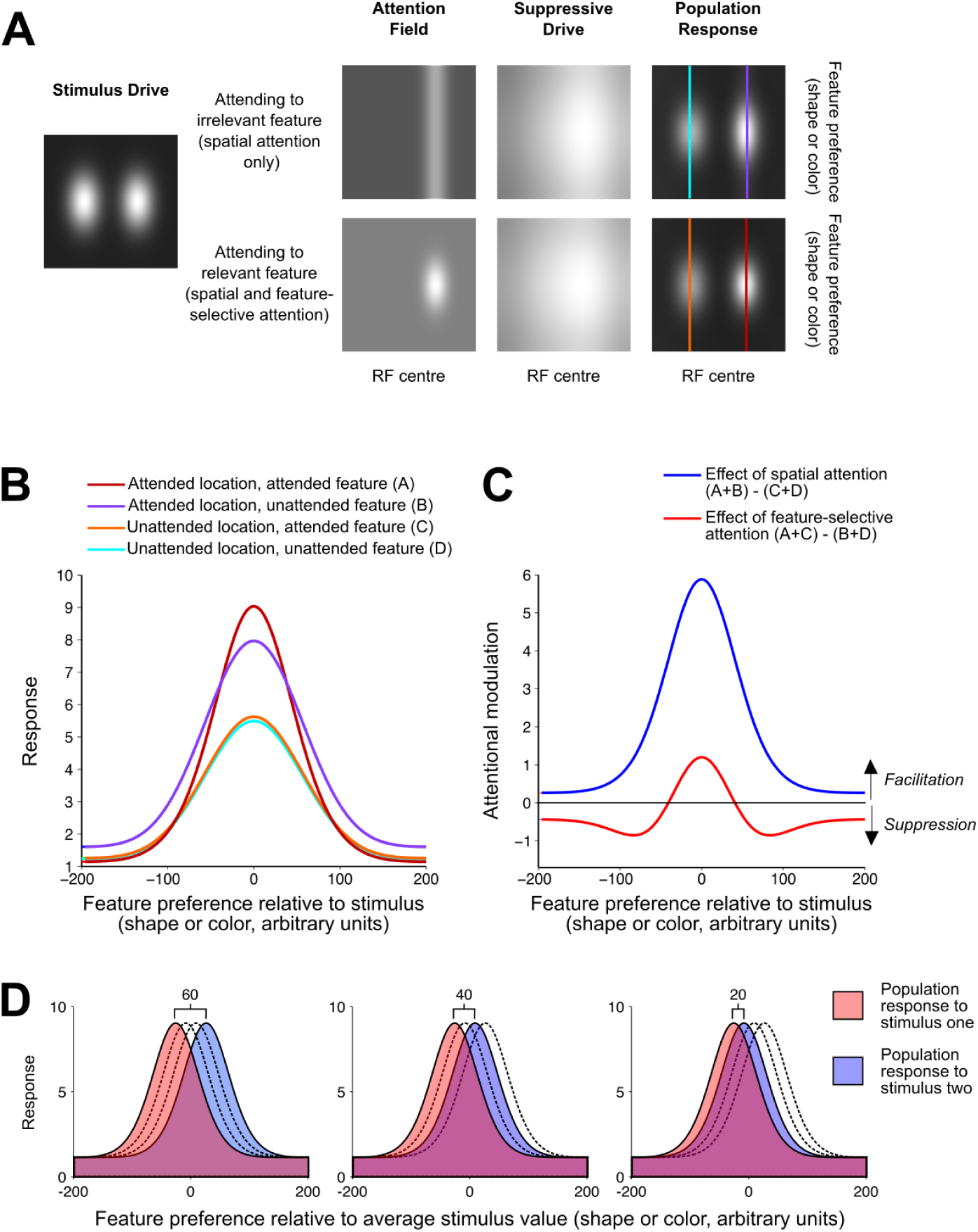
Summary of a normalization model of attention (Reynolds and Heeger, 2009), as implemented here to predict the effects of spatial and feature-selective attention on classifier performance. **A**: an illustration of each of the model elements from Reynolds and Heeger (2009), Figure 1, for a set of example model parameters, where each grayscale image depicts a matrix of values varying along a spatial dimension (horizontally) and a feature dimension (vertically). For each set of model parameters we generated a single ‘stimulus drive’, and two versions of the ‘attention field’, which lead to subtly different ‘suppressive drives’ and ‘population responses’. From these two population responses we derived curves predicting the population response as a function of each neuron’s preferred feature value for each of the four attention conditions (the columns of the matrix indicated with different colored vertical lines in **A**). These population responses are plotted again as lineplots in **B**. In **C** (redrawn from Figure 6A) the predicted effects of spatial and feature-based attention on the population response are summarized as the difference between relevant population curves from **B**. **D:** We predicted classifier performance in each attention condition by centering the population response from **B** on 4 different stimulus feature values and predicting classifier performance when discriminating between population responses to stimuli of that were either 60, 40 or 20 (arbitrary) units apart along the feature dimension, to simulate the population response to stimuli that were 3, 2 or 1 steps apart in either color or shape. We predicted classifier performance (d’) using the separation of the two population responses, in a manner analogous to that used in signal detection theory (see Supplementary Methods for details)

### MEG data analysis: Granger analysis of feedforward and feedback information flows

We tested for temporal dependence between the patterns of classifier performance in occipital and frontal datasets, seeking evidence of information flows from occipital to frontal cortices (feedforward) and from frontal to occipital cortices (feedback), following the rationale we developed in earlier work (Goddard et al., 2016). Specifically, we tested for Granger causal relationships between the patterns of classifier performance based on the occipital and frontal datasets. We summarized the color and shape information for each region (occipital and frontal), at each timepoint, as a 6×4 dissimilarity matrix (DSM) of classifier performances. For both color and shape, the 6×4 DSM was defined as each pairwise comparison (6 classifications across the 4 levels of the feature), by 4 attention conditions (aLaF, aLuF, uLaF, uLuF).

The logic of Granger causality is that time series X ‘Granger causes’ time series Y if X contains information that helps predict the future of Y better than information in the past of Y alone (for a recent review of its application in neuroscience, see Friston et al. (2013)). We performed a sliding-window analysis of a simplified (special case) of Granger causality, using the partial correlations in Equations 3 and 4 to define ‘Feedforward’ (*FF*) and ‘Feedback’ (*FB*) information flows at each time point (*t*).

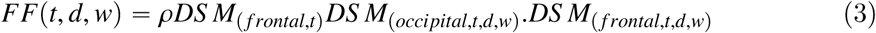

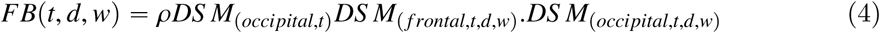

where *DS M*_(*loc*,*t*)_ is the DSM based on the sources at location *loc* at time *tms* post stimulus onset, and *DS M*_(*loc*,*t*,*d*,*w*)_ is the DSM based on the sensors at location *loc*, averaged across all time points from *t*–*dms* to *t*–(*d* + *w*)*ms* post stimulus onset. We calculated *FF* and *FB* for 30 overlapping windows: for 5 window widths (*w* = 10, 20, 30, 40 or 50 ms) for each of 6 delays (*d* = 50, 60, 70, 80, 90 or 100). We tried a range of values for *w* and *d* in order to capture interactions between occipital and frontal cortices that may occur at different timescales.Since the results were broadly similar across values of *w* and *d* (see Figure S8) we report *FF* and *FB* values averaged across all values of *w* and *d*.

We report the results of this analysis in terms of the difference between the feedforward and feedback information flows (FF-FB). To assess whether this difference was significantly above or below chance, we generated a null distribution of this difference at every timepoint by performing the same analysis on 1000 bootstraps of data from each subject where the exemplar labels were randomly permuted for each of the DSMs used in Equations 3 and 4.

### Data availability

All the raw data and the results of our classification analyses are available on an Open Science Framework project (after publication we will make this project publically accessible and include the DOI for the project in our manuscript).

## Acknowledgments

This project was funded under an Australian Research Council Future Fellowship (FT120100816) awarded to TC, ARC Discovery Projects (DP160101300) awarded to TC and (DP170101840) awarded to AW, an ARC Future Fellowship (FT170100105) awarded to AW, MRC (U.K) intramural funding SUAG/035/RG91365 awarded to AW, and the ARC Centre of Excellence in Cognition and its Disorders (CE110001021). We thank Erika Contini and Elizabeth Magdas for their assistance with MEG data collection.

## Supplementary Material

### Supplementary 1: Event related potentials

**Figure S1:**
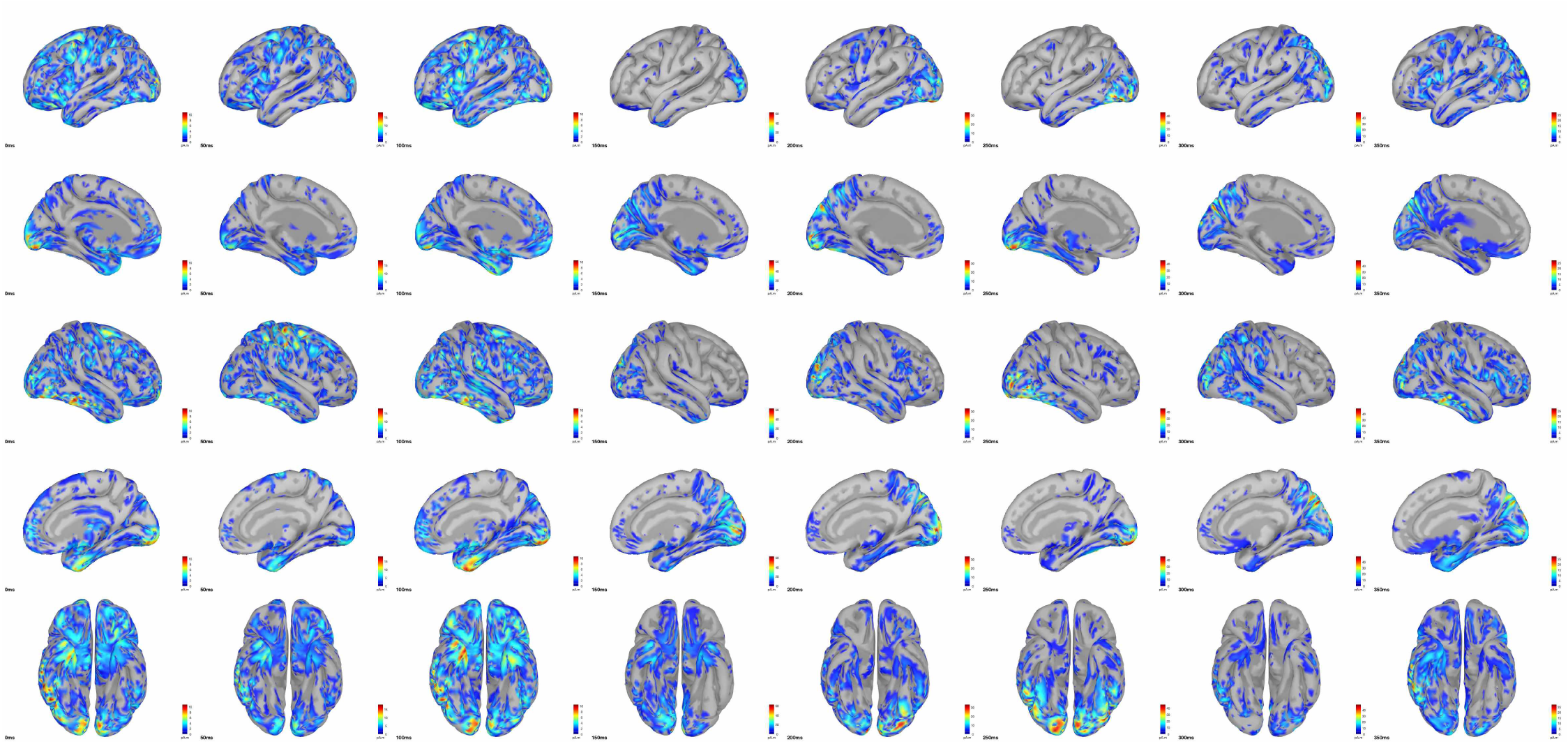
Event related potentials. Here the average event related potentials (ERPs), across all 2048 trials, are shown (averaged across 20 subjects), for 8 time points evenly spaced between 0ms and 350ms after stimulus onset. Each column shows the same 5 views of the brain at a single time point, and the ERPs are thresholded separately for each column so that only those values that exceed 10% of the maximum ERP at that time point are shown. The peak ERP values across these time points were found from 150–250*ms* after stimulus onset, and the potentials were of greatest amplitude at this time were around the occipital pole and surrounding cortex, consistent with a visually evoked response.

### Supplementary 2: Modelling: extended methods and results

We started with the Matlab routines from Reynolds and Heeger (2009) that are freely available from http://www.cns.nyu.edu/heegerlab/. Since we did not have strong a priori predictions for many of the model parameters, we tested a broad range of plausible model parameters (see Table 1). For each set of model parameters (172,800 sets in total) we used the Reynolds and Heeger (2009) model to predict the response of the neural population as a function of stimulus feature preference (along the shape or color dimension), for each of four cases, illustrated by lines of different colors in Figure 7**A-B**. In every case the stimulus was a single feature value (a specific color or shape) at 2 fixed locations (left and right of fixation). In two cases, we simulated attention to one location in the absence of any feature-based attention (simulating attention to the orthogonal feature dimension). In the other two cases we simulated attention to one location and attention to the feature value of the stimuli. From these we predicted the population response at attended and unattended locations, in the presence and absence of feature-based attention. As illustrated in Figure 7**C**, according to the model spatial attention tends to boost the population response as a multiplicative scaling of the original response, while feature-based attention produces both facilitation and suppression of the response which leads to sharpening of the population response around the attended value.

**Table 1:** Model parameters from the normalization model of attention (Reynolds and Heeger, 2009) that we used in model simulations. We defined the stimulus and response matrices as varying from −200 to 200 along both spatial and feature dimensions (arbitrary units). We generated the model predictions for every combination of the above model parameters, resulting in 172,800 sets of model predictions. The process of estimating classifier accuracy from the model predictions is summarized in Figure 7.

| Model parameter | Parameter description | Values tested |
| --- | --- | --- |
| *stimWidth* | Spatial extent of stimulus | 25 (Fixed value) |
| *stimFeatureWidth* | Extent of stimulus along feature dimension | 25 (Fixed value) |
| *ExWidth* | Spread of stimulation field along spatial dimension | 30, 40, 50, 60, 70, 80, 90 or 100 |
| *EthetaWidth* | Spread of stimulation field along feature dimension | 30, 40, 50, 60, 70 or 80 |
| *IxWidth* | Spread of suppressive field along spatial dimension | $=C*ExWidth$, where $C=1.5, 2$ or $2.5$ |
| *IthetaWidth* | Spread of suppressive field along feature dimension | $=C*EthetaWidth$, where $C=1.5, 2$ or $2.5$ |
| *AxWidth* | Extent/width of the spatial attention field | $=ExWidth$ |
| *AthetaWidth* | Extent/width of the featural attention field | $=EthetaWidth$ |
| *ApeakX* | Peak amplitude of spatial attention field | 2, 4, 6 or 8 |
| *ApeakTheta* | Peak amplitude of the feature-based attention field | 2, 4, 6 or 8 |
| *Abase* | Baseline of attention field for unattended locations/features | 1 (Fixed value) |
| *baselineMod* | Amount of baseline added to stimulus drive | 0, .1, .3, .5 or 1 |
| *baselineUnmod* | Amount of baseline added after normalization | 0, .1, .3, .5 or 1 |
| *sigma* | Constant that determines the semi-saturation contrast | $1e-6$ (Fixed value) |
| *Ashape* | either ‘oval’ or ‘cross’ | ‘oval’ (Fixed value) |

One difference between the Reynolds and Heeger (2009) model and our experiment is that the model is designed to capture feature-based attention (attending to a specific feature value, e.g. red), whereas we manipulated feature-selective attention (attending to a feature dimension, e.g. color). While feature-based attention has received greater attention in the electrophysiology literature, feature-selective attention has been demonstrated to have similar effects at the level of single neurons (Cohen and Maunsell, 2011). We therefore implemented the feature-selective attention manipulation in the model by generating population responses to two stimuli of the same feature value, and modeling the presence of feature-selective attention as feature-based attention to that feature value.

For every predicted population response we predicted classifier performance when discriminating responses to stimuli of different feature values. To do this we compared two population responses that were identical except that they were centered on different feature values, as shown in Figure 7**D**. To simulate the three steps of stimulus difference, we considered cases where the centers of the population responses were separated by either 20, 40 or 60 in the arbitrary units of the feature dimension. In the case of stimuli varying in color, the chromaticity coordinates of the stimuli varied from strongly red u’v’: 0.35, 0.53, to strongly green u’v’: 0.16, 0.56, which means that for the model we were treating a difference of 60 arbitrary units as a distance of approximately 0.19 in the u’v’ chromaticity plane. For shape the feature dimension is defined by the transition from ‘X-shaped’ to ‘non-X-shaped’. We are not asserting that there exist neurons tuned to this novel complex shape dimension in the same way as there are neurons tuned to color, but for the purposes of the model we treated these dimensions as equivalent. Since subject performance was similar for the color and shape task, we used the same distances (20, 40 and 60 in the arbitrary units) to avoid adding another parameter to the modeling results.

Using the pairs of population responses (such as those in Figure 7**D**) we predicted classifier performance (d’) using the separation of the two population responses, in a manner analogous to that used in signal detection theory. To determine d’ for these population responses we calculated a ‘hit rate’ for an optimal observer detecting a signal (stimulus two) amongst noise (stimulus one), where their criterion (*c*) is at the midpoint between the peaks of the two curves. We defined the ‘hit rate’ (*hits*) as the area under the blue curve to the right *c*, and the ‘false alarm rate’ (*FA*) as the area under the red curve to the right of *c*. Then the predicted classifier performance d’ = norminv(hits) - norminv(FA). In this way, for each set of model parameters we predicted classifier performance in each attention condition, for each of the three step sizes in feature difference.

From the predicted classification performance, we summarized the predicted effects of spatial attention and feature-selective attention using the **SpatAtt** and **FeatAtt** values from equations 1 and 2.

**Figure S2:**
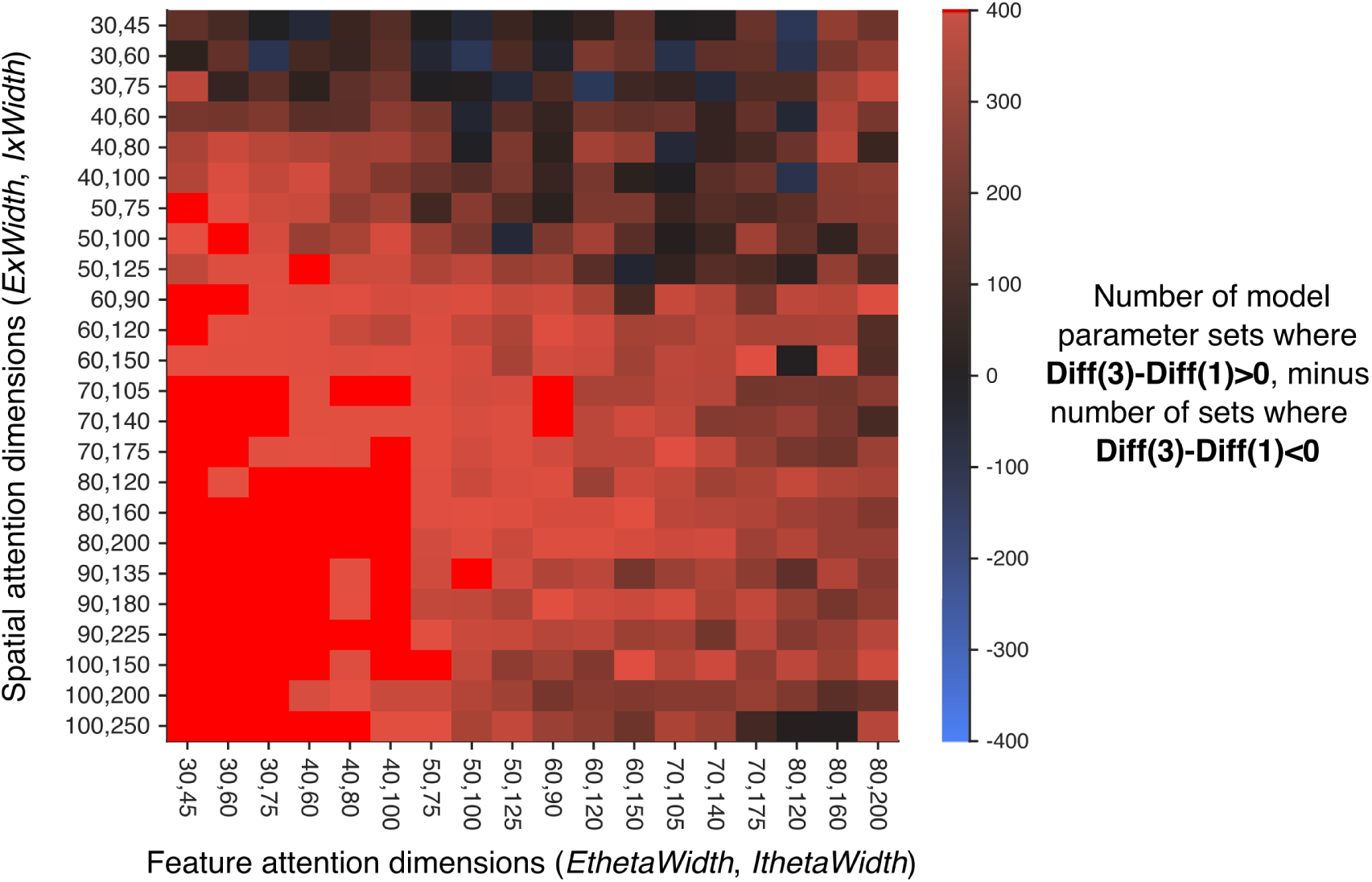
Comparing the model predictions across 4 model parameters. The model predictions across 4 model parameters: the excitation and inhibition width of the spatial and feature-based attention fields (*ExWidth*, *IxWidth*, *EthetaWidth* and *IthetaWidth* in Table 1). In each cell, there were 400 sets of model parameters (where other model parameters were varied). For each set of model parameters, we calculated the difference between attention effects (**Diff = SpatAtt-FeatAtt**) across feature difference (as in Figure 6). Here we show number of model parameter sets for which the pattern of results was qualitatively similar to the average model prediction (Figure 6**B**) and to the data (e.g. Figure 6**E**). That is, model sets where **Diff** at 3 steps (**Diff(3)**) minus **Diff** at 1 step difference (**Diff(1)**) was positive (red cells, 95% of cases). There were also some combinations of excitation and inhibition widths for which all 400 cases followed this pattern (bright red cells, 16% of cases).

### Supplementary 3: Control analysis on the effects of spatial bias in fixation location

To encourage participants to suppress eye movements we provided explicit instructions to maintain fixation on the constantly-present fixation cross, and we informed participants that we were using an eye tracker to measure their eye movements. We also informed participants that the onset of each trial was contingent on the eye tracker detecting their fixation. We chose a short stimulus duration (maximum 150*ms*) to discourage eye movements after the onset of the stimulus, and if the eye tracker indicated the participant was no longer fixating the stimulus was removed immediately.

Due to eye tracker variability we treated fixation locations within 1 dva of the center of the screen as ‘fixating’ for the purposes of the fixation-contingent onset, in order to avoid extensive delays in the experiment. Because of this, we could not exclude the possibility that participants had a small bias to fixate slightly towards the attended location. From the eye tracking data, we found that most participants (16 of 20) showed a small bias to fixate slightly towards the attended location (see Figure S3). To check that this bias was not driving the differences we observed between spatial and feature-selective attention, we repeated our group analyses including only the 4 participants who had a small bias to fixate towards the unattended location, and found the same pattern of results as in the main analyses (Figure S4).

**Figure S3:**
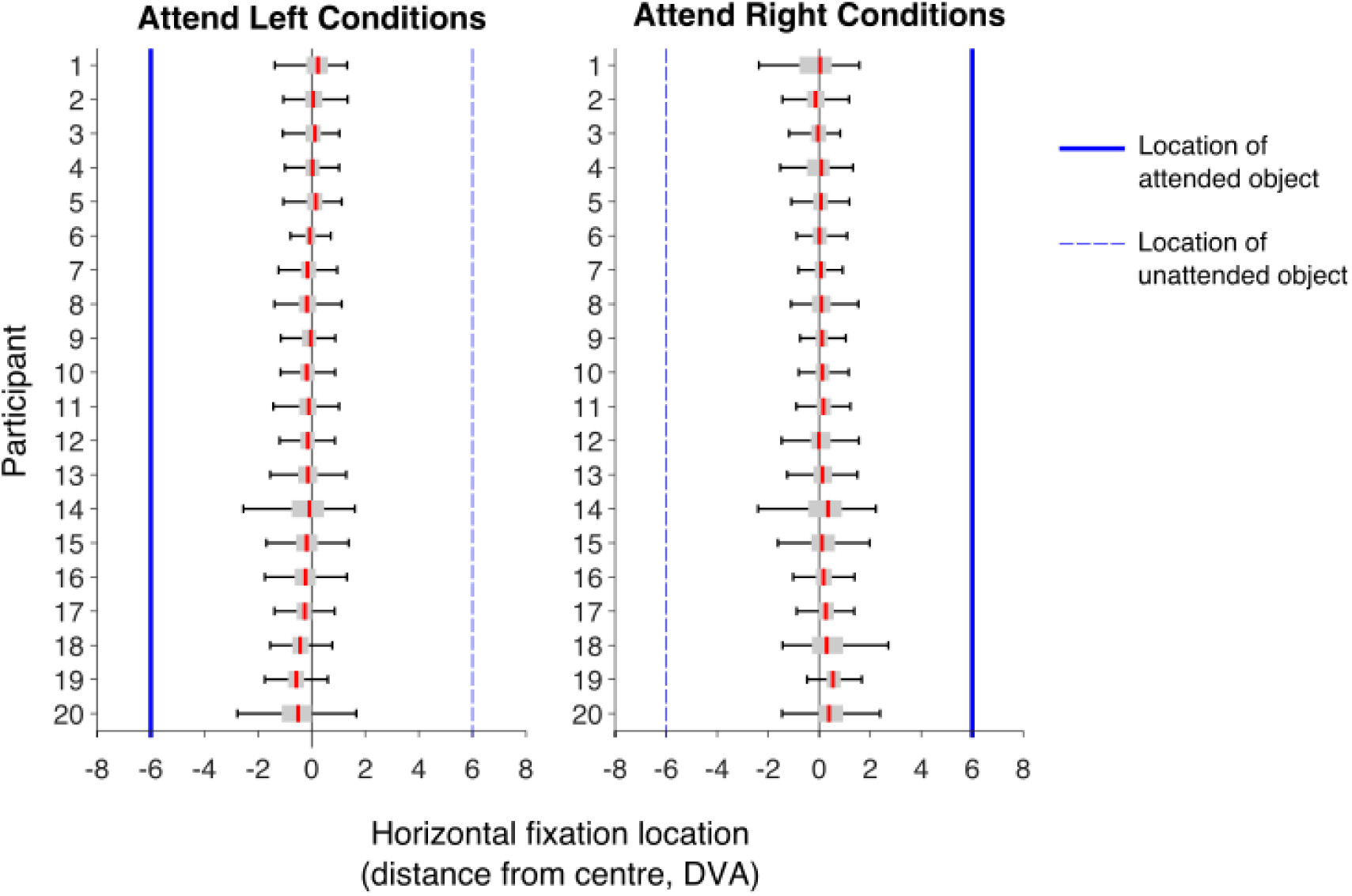
Distributions of fixation locations, for individual participants. In each distribution, red lines show the median, and the shaded gray box indicates the first and third quartiles of the distribution of 1024 fixation locations. Participants are ordered by their overall bias, from biased towards unattended to biased towards attended location.

**Figure S4:**
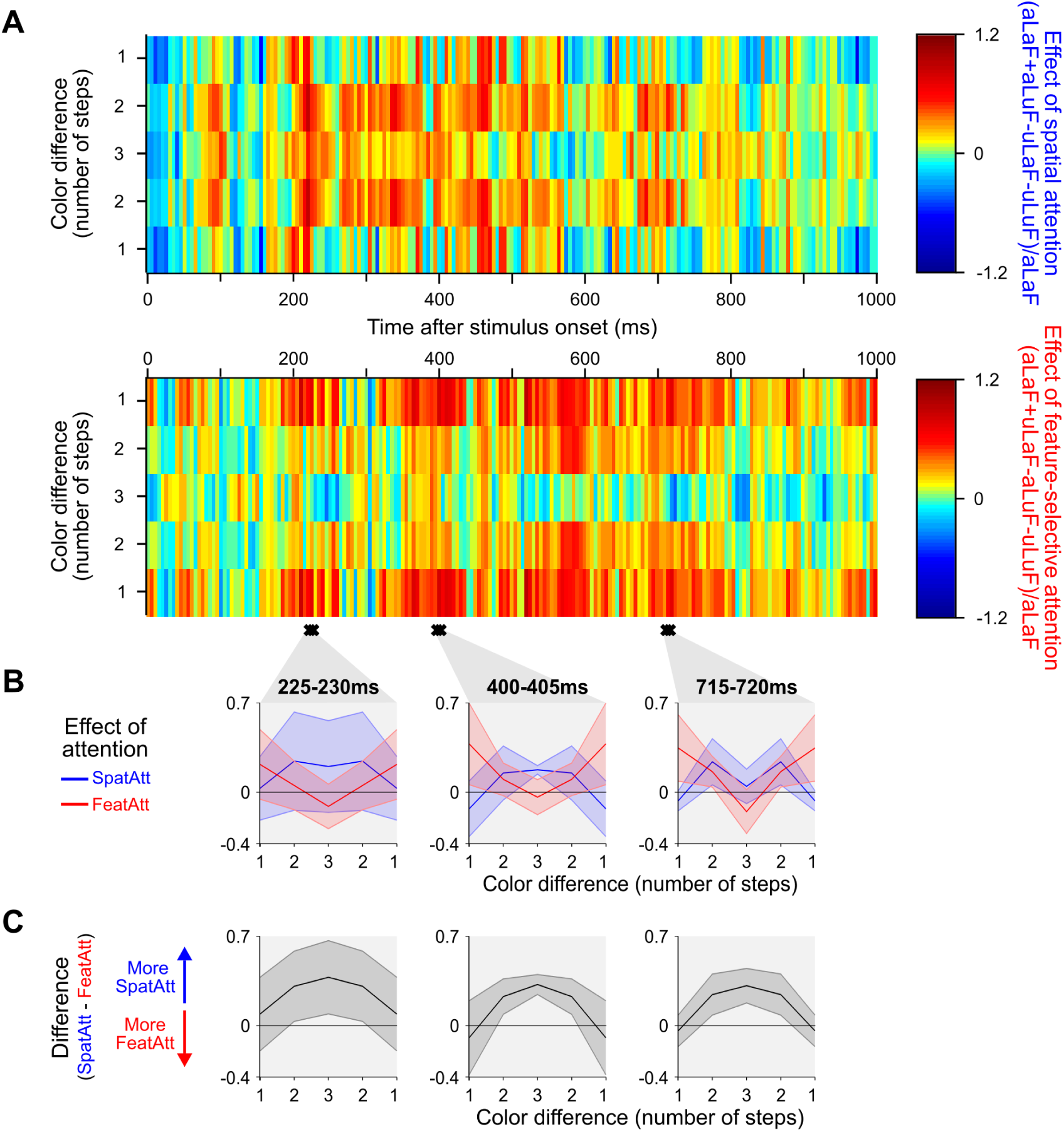
Effects of spatial and feature-selective attention across decoding of object color in occipital ROIs for participants with a slight bias to fixate toward the unattended location. Results for a subset of participants (n=4, participants 1-4 in Figure S3). Plotting conventions for **A**-**C** are as in Figure 6**C-E**.

### Supplementary 4: Effects of spatial and feature-based attention on decoding of shape

**Figure S5:**
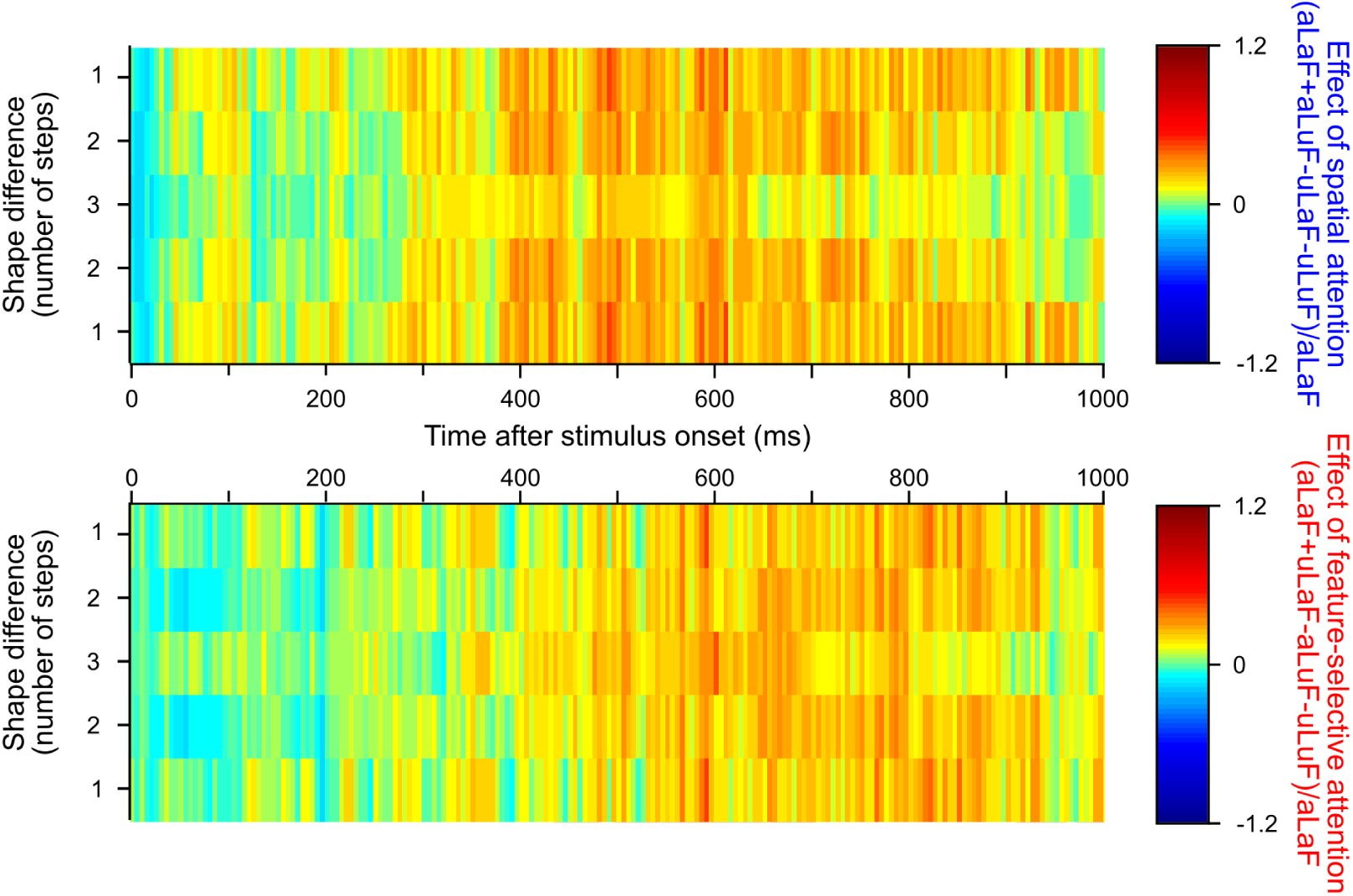
Effect of spatial and feature-based attention on the decoding of object shape in the occipital ROI. Plotting conventions as in Figure 6**C**. In this case, there were no consecutive time points at which there was a significant interaction between attention type and step size.

**Figure S6:**
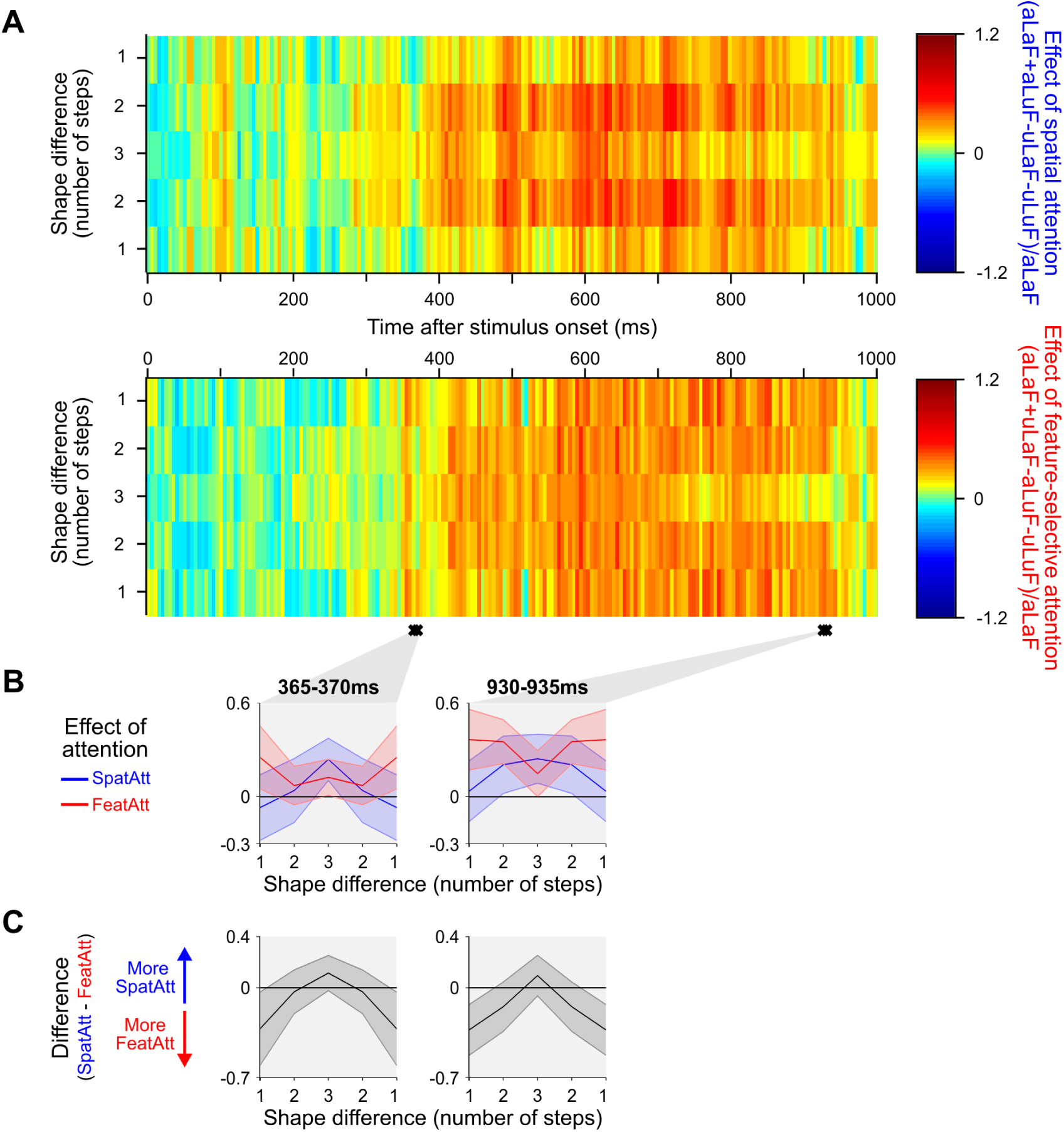
Effects of spatial and feature-selective attention across decoding of object shape for all MEG sensors. Plotting conventions for **A**-**C** are as in Figure 6**C-E**.

### Supplementary 5: Information flow analysis, varying averaging window

**Figure S7:**
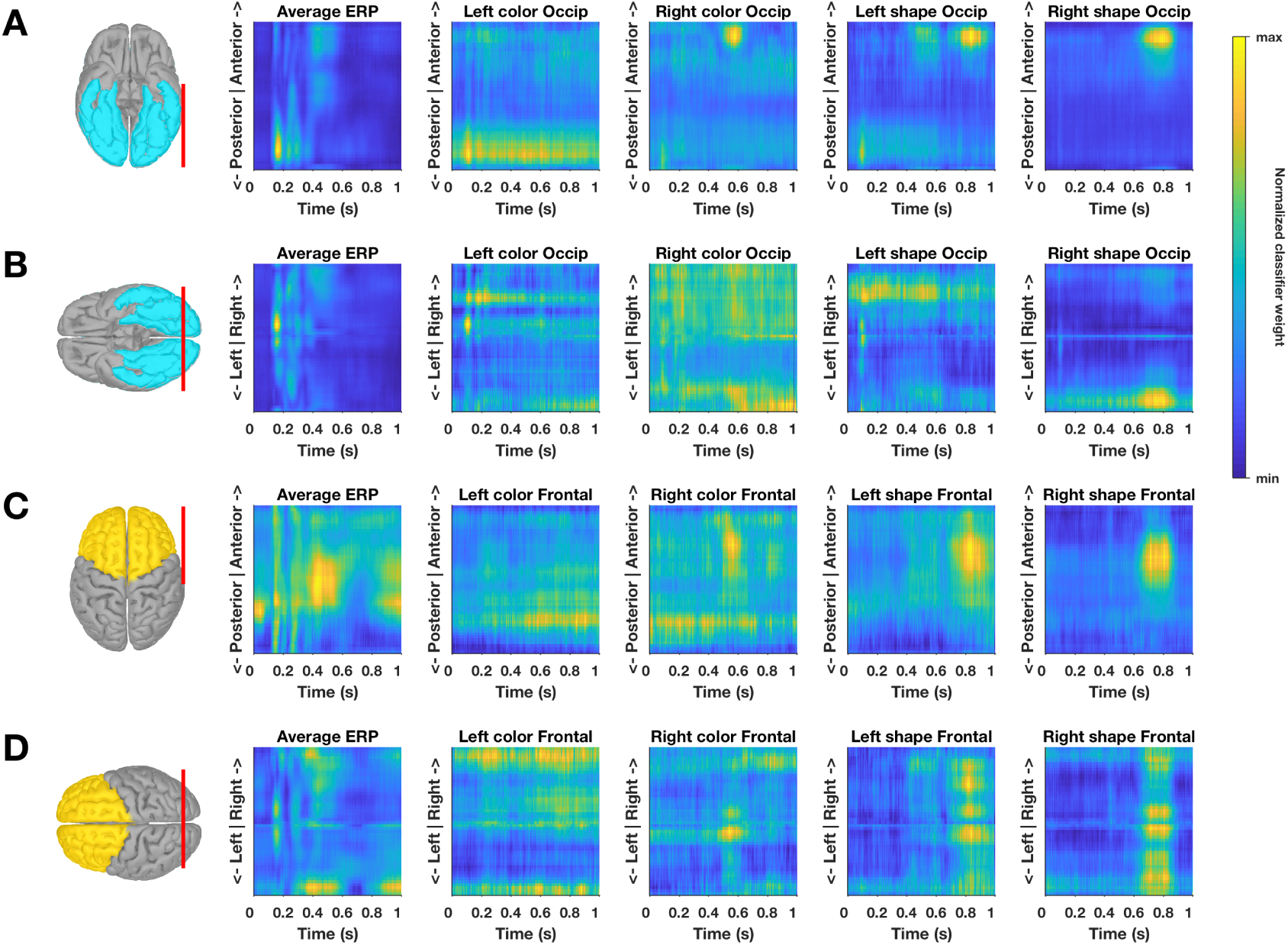
Average event-related potential (ERP) and transformed classification weights (W’). For both the occipital (**A-B**) and frontal (**C-D**) ROIs we spatially binned the ROI into 50 equally spaced bins across two dimensions: posterior to anterior (**A,C**) and left to right (**B,D**). In each subplot, the bins span the distance indicated by the red line over the ROI in the leftmost column. In the remaining columns, we plot, as a function of time, the average ERP (2nd column) and average transformed classifier weights (**W’**, see methods) for decoding in the attended location, attended feature condition (columns 3-6). That is, ‘Left color’ (column 3) is the decoding of the color of the left object color when performing the color task on the left object, ‘Right shape’ (column 6) is the decoding of the shape of the right object shape when performing the shape task on the right object, etc. The occipital ROI showed a lateralization consistent with classifier performance being driven by retinotopically organized visual cortex: when decoding of features of the stimulus in the left visual field the classifier tended to give higher weight to right hemisphere locations, and vice versa. The frontal ROI did not show clear evidence of lateralization, consistent with frontal regions containing information about both contra- and ipsilateral visual fields (e.g. Lennert and Martinez-Trujillo (2013)).

### Supplementary 6: Information flow analysis, varying averaging window

**Figure S8:**
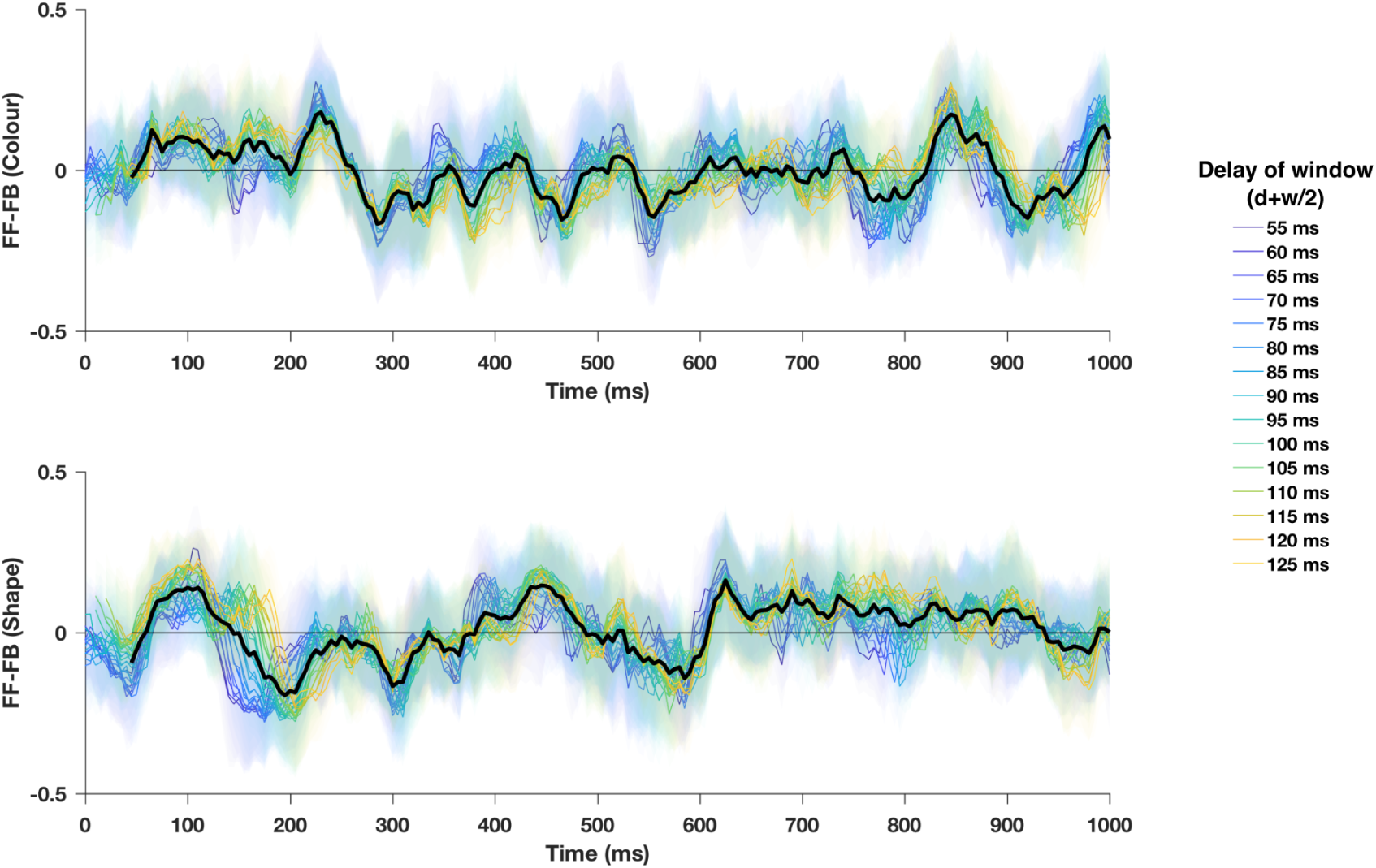
Information flow analysis across varying averaging windows. Upper and lower plots show, for color and shape respectively, the direction of information flow (FF-FB) for each averaging window, given by 5 window widths (*w* = 10, 20, 30, 40 or 50 ms) for each of 6 delays (*d* = 50, 60, 70, 80, 90 or 100). Lines are colored according to the midpoint of the window, and translucent shaded error bars of the same colour indicate the 95% confidence intervals of each between-subject mean. The thick black line shows the average of these lines, replotted from Figure 5**A** (see Figure 5**A** for confidence intervals of this average).

## Footnotes

1 In Reynolds and Heeger (2009) the parameter ‘Ashape’ was set to ‘oval’ rather than ‘cross’ for all but one of their figures, but to our knowledge our result is the first test of this prediction.

